# The genetic architecture of fitness drives population viability during rapid environmental change

**DOI:** 10.1101/660803

**Authors:** Marty Kardos, Northwest Fisheries Science Center, Gordon Luikart

## Abstract

The rapid global loss of biodiversity calls for improved predictions of how populations will evolve and respond demographically to ongoing environmental change. The heritability (*h*^2^) of selected traits has long been known to affect evolutionary and demographic responses to environmental change. However, effects of the genetic architecture underlying the *h*^2^ of a selected trait on population responses to selection are less well understood. We use deterministic models and stochastic simulations to show that the genetic architecture underlying *h*^2^ can dramatically affect population viability during environmental change. Polygenic trait architectures (many loci, each with a small phenotypic effect) conferred higher population viability than genetic architectures with the same initial *h*^2^ and large-effect loci under a wide range of scenarios. Population viability also depended strongly on the initial frequency of large-effect beneficial alleles, with moderately low initial allele frequencies conferring higher viability than rare or already-frequent large-effect alleles. Greater population viability associated with polygenic architectures appears to be due to higher short term evolutionary potential compared to architectures with large-effect loci. These results suggest that integrating information on the trait genetic architecture into quantitiative genetic and population viability analysis will substantially improve our understanding and prediction of evolutionary and demographic responses following environmental change.

## Introduction

One of the most urgent undertakings for science is to understand how biodiversity will respond to human-driven environmental change (Mills et al. 2018; Nadeau and Urban 2019; Stockwell et al. 2003; Urban et al. 2016; Wilson 2016). Populations can persist through environmental change either by shifting their geographic distributions to track suitable habitats, or by adapting to changing local conditions (Pease *et al.* 1989). Predicting how populations will evolve and respond demographically to selection imposed by environmental change (e.g., global warming) is a difficult task, but crucial to understanding and mitigating the ongoing extinction crisis (Alberto et al. 2013; Chevin and Lande 2010; Funk et al. 2018; Shaw 2019; Stockwell et al. 2003; Urban et al. 2016).

This need has motivated several theoretical and simulation-based analyses of evolutionary and demographic responses to selection induced by environmental change (Bay et al. 2017; Gomulkiewicz et al. 2010; Lynch et al. 1991; Nunney 2015; Pease et al. 1989). These studies generally combined models of the genetic basis of a selected phenotype, fitness as a function of phenotype, and density-dependent fitness to link adaptation to population dynamics under environmental change. Such models can be used to identify at-risk populations, and to identify the factors that most strongly affect population responses to environmental change and potential resource management strategies to mitigate extinction risk.

Realistic genetic models of variation in selected phenotypes are crucial for inferring evolutionary and demographic responses to selection. The expected phenotypic response per generation has long been known to be proportional to the selected trait’s heritability (*h*^2^, the proportion of phenotypic variance due to additive genetic effects). *h*^2^ is therefore a key genetic parameter for modelling evolutionary and demographic responses to environmental change (Chevin and Lande 2010; Falconer and Mackay 1996; Gomulkiewicz and Holt 1995; Lynch and Lande 1993; Nadeau and Urban 2019; Urban et al. 2016). Population genetics theory shows that the genetic architecture of a trait (i.e., the number, distribution of effect sizes, and allele frequencies of the loci underlying *h*^2^) can strongly affect the temporal dynamics of *h*^2^ and set the limits of adaptive phenotype evolution (Chevalet 1994; Walsh and Lynch 2018).

Polygenic traits (affected by many loci, each with a small effect) are expected to have higher evolutionary potential than traits with large-effect loci and the same initial *h*^2^. This is because *h*^2^ and the rate of adaptation are expected to decline more rapidly during adaptation for traits with large-effect loci than when a selected trait is polygenic (Barton and Keightley 2002; Chevalet 1994; Walsh and Lynch 2018). This makes the scope for potential adaptive phenotypic evolution generally larger for polygenic traits than for traits with the same initial *h*^2^ and large-effect loci. Populations with polygenic selected phenotypes may therefore be substantially more likely to adapt to new conditions, and to remain viable through environmentally-induced selection than when large-effect loci are responsible for much of the *h*^2^. Knowing the initial *h*^2^ of the selected trait, and using realistic models of the genetic basis of phenotypic variation could hence be crucial to inferring biological responses to environmental change. However, most previous analyses of the response to environmental change either didn’t measure *h*^2^, or assumed that *h*^2^ was constant during bouts of selection (Bay et al. 2017; Bürger and Lynch 1995; Chevin 2019; Gomulkiewicz and Holt 1995; Gomulkiewicz et al. 2010; Lande 1983; Lynch and Lande 1993; Nunney 2015; Pease et al. 1989). For example, Nunney (2015) and Bay et al. (2017) did not account for *h*^2^ in their population genetic analyses of the effects of trait genetic architecture on popultion dynamics. Lande (1983), Gomulkiewicz et al. (2010), and Chevin (2019) assumed that the genetic variance contributed by loci with small effects remained constant through time. However, the selection response is expected to alter the genetic variance, and the resulting temporal variation in *h*^2^ can substantially affect the evolutionary response (Walsh and Lynch 2018).

Omitting *h*^2^ or the effects of genetic architecture on temporal variation in *h*^2^ may result in unreliable inferences of evolutionary and demographic responses to environmental change. Recent studies show that many fitness-related traits are highly polygenic (Boyle et al. 2017). Assuming that *h*^2^ is constant through time despite adaptive evolution – consistent with the infinitesimal model of inheritance in a large population – may be reasonable in such cases. Many other traits, including some that are likely important for adaptation to climate change (Thompson et al. 2019), are governed by loci with very large phenotypic effects and a broad range of allele frequencies (Barson et al. 2015; Epstein et al. 2016; Jones et al. 2018; Kardos et al. 2015; Lamichhaney et al. 2016; Pearse et al. 2019; Thompson et al. 2019). This emerging picture of a large diversity in the genetic architecture of fitness traits, and the importance of genetic architecture to adaptive potential, suggests that including information on both the initial *h*^2^ and the underlying genetic architecture of the selected phenotype(s) might substantially improve our understanding and prediction of evolutionary and demographic responses to environmental change.

The objective of this paper is to determine when the genetic architecture of a selected phenotype affects the viability of populations subjected to a shifting phenotypic optimum caused by environmental change. To address this, we developed deterministic evolutionary-demographic models, and stochastic, individual-based simulations that account for the initial *h*^2^ and the effects of the genetic architecture on temporal change of *h*^2^.

## METHODS

### A deterministic model of population responses to environmental change

We first develop a deterministic, evolutionary-demographic model that builds upon previous approaches used to investigate evolutionary rescue (Chevin and Lande 2010; Gomulkiewicz and Holt 1995; Gomulkiewicz et al. 2010; Lande 1983; Lynch and Lande 1993). We use this model to determine expectations for phenotypic evolution and population growth under a range of simple genetic architectures with purely additive phenotypic effects, multiple unlinked loci with equal phenotypic effects, and no linkage disequilibrium, epistasis or plasticity. Further down we evaluate the effects of linkage disequilibrium, and varying phenotypic effects among loci in the analysis of this model.

We model sexually reproducing, non-selfing, diploid populations that have discrete generations and follow a discrete logistic model of density-dependent population growth (May 1974). Individual fitness is a Gaussian function of a quantitative trait, with the fitness of an individual with phenotype value *z* being

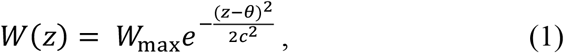

where *W*_max_ is the fitness (expected lifetime reproductive success) of an individual with optimum phenotype value *θ* when the population size *N* is very close to zero, and *c*^2^ defines the width of the fitness function. The population has an initial mean phenotype of 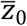 equal to the initial phenotypic optimum *θ*_0_. The selected phenotype is assumed to be normally distributed with additive genetic (*V*_G_) and random environmental (*V*_E_) variance components summing to the total phenotypic variance *V*_z_ (*h*^2^ = *V*_G_/*V*_z_). The phenotype’s probability density function is

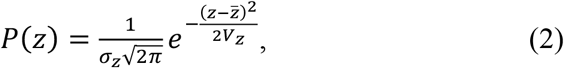

Where 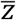 is the mean phenotype, and σ_*z*_ the phenotype standard deviation. 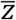 is calculated as

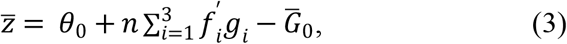

where *f′*_*i*_ is the frequency of the *i*^th^ of the three possible genotypes per di-allelic locus, *g*_*i*_ is the genetic value of the *i*^th^ of the three possible genotypes per locus, *n* is the number of diallelic loci affecting the trait, and 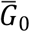 is the value of the second term (i.e., the mean additive genetic value among individuals in the population) in the first generation. *g*_*i*_ is calculated as

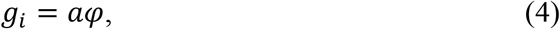

where *a* is half the phenotypic difference between the two alternative homozygous genotypes, and *φ* is the number of copies of the allele that confers a larger phenotype (the A1 allele) in the ith of the three possible genotypes. The third term in (3) ensures that 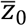 is exactly equal to *θ*_0_, and is necessary because the focal allele at each locus increases the size of the phenotype (i.e., *a* in (4) is always positive). A sudden environmental change permanently shifts *θ* from its initial value *θ*_0_ in the first generation to *θ*_1_, thus imposing directional selection on the phenotype and an environmental challenge to population persistence.

We assume that the A1 allele has the same initial frequency *p*_0_ at each locus. Further, the frequency of the A1 allele(s) is assumed to evolve identically at each of the *n* loci, such that *p* in generation *t* + 1 at each locus is

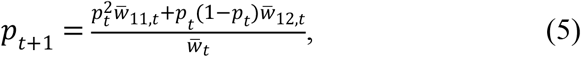

where 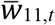 and 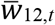 represent the mean relative fitness of homozygous A1A1 genotypes, and heterozygous A1A2 genotypes in generation *t*, respectively, and 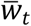is the mean individual fitness in generation *t*. Mean absolute individual fitness in the population is calculated by integrating over the product of the fitness and phenotype density functions:

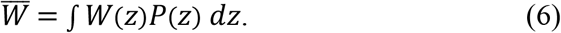

The mean genotype-specific relative fitness (i.e., 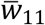 or 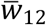) is calculated as in (5) except with the variance (*V*_z_) and mean 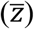 of the phenotype probability density function in (2) being conditional on holding the genotype constant at a locus. The *V*_z_ conditional on holding the genotype constant at a locus is

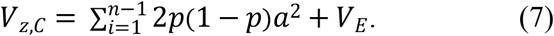

The mean phenotype conditional on holding the genotype constant at a locus is

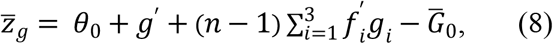

where *g*′ is the genetic value of the single-locus genotype being held constant (i.e., *g*′ = 0 for genotype A2A2, *g*′ = *a* for A1A2, and *g*′ = 2*a* for A1A1).

We calculate *h*^2^ each generation as

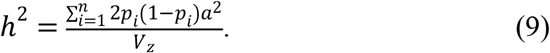

Population size (*N*) in generation *t* + 1 is calculated following the discrete logistic model as

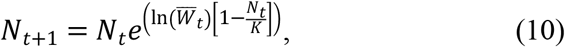

where *K* is the carrying capacity. We numerically iterated this model for 80 generations to evaluate the effects of the number of loci underlying *h*^2^ on the evolutionary and demographic responses to a sudden shift in the optimum phenotype due to an environmental change.

We chose combinations of parameter values to test effects of the genetic architecture of a relatively highly heritable trait on population persistence under strong environmentally-induced selection (i.e., *θ*_1_ in the far right tail of the initial phenotype distribution, Figure 1). We first considered the simple case where either 1 or 2 large-effect loci (large-effect architectures), or 100 loci with small effects (polygenic architecture) contributed to *V*_G_. We set parameters values as maximum fitness *W*_max_ = 1.5, initial heritability 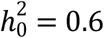, initial phenotype variance to *V*_P_ = 10, initial mean phenotype 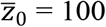, initial optimum phenotype *θ*_0_ = 100 (in arbitrary units), new optimum phenotype *θ*_1_ = 110 (3.2 standard deviations from *θ*_0_), width of the fitness function *c* = 6, the initial population size *N*_0_ = 500, carrying capacity *K* = 1,000. Each of the small-effect loci contributed equally to *V*_G_. The fitness function and the initial phenotype probability density distribution are shown in Figure 1. This combination of parameters yields an initial mean absolute fitness of 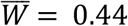 and threfore a rapid initial decline in population size. We considered a population extinct when *N* was < 2. Note that this model, and the models below, control for the initial evolvability (mean-scaled additive genetic variance) (Hansen et al. 2011) in addition to 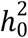.

**Figure 1.**
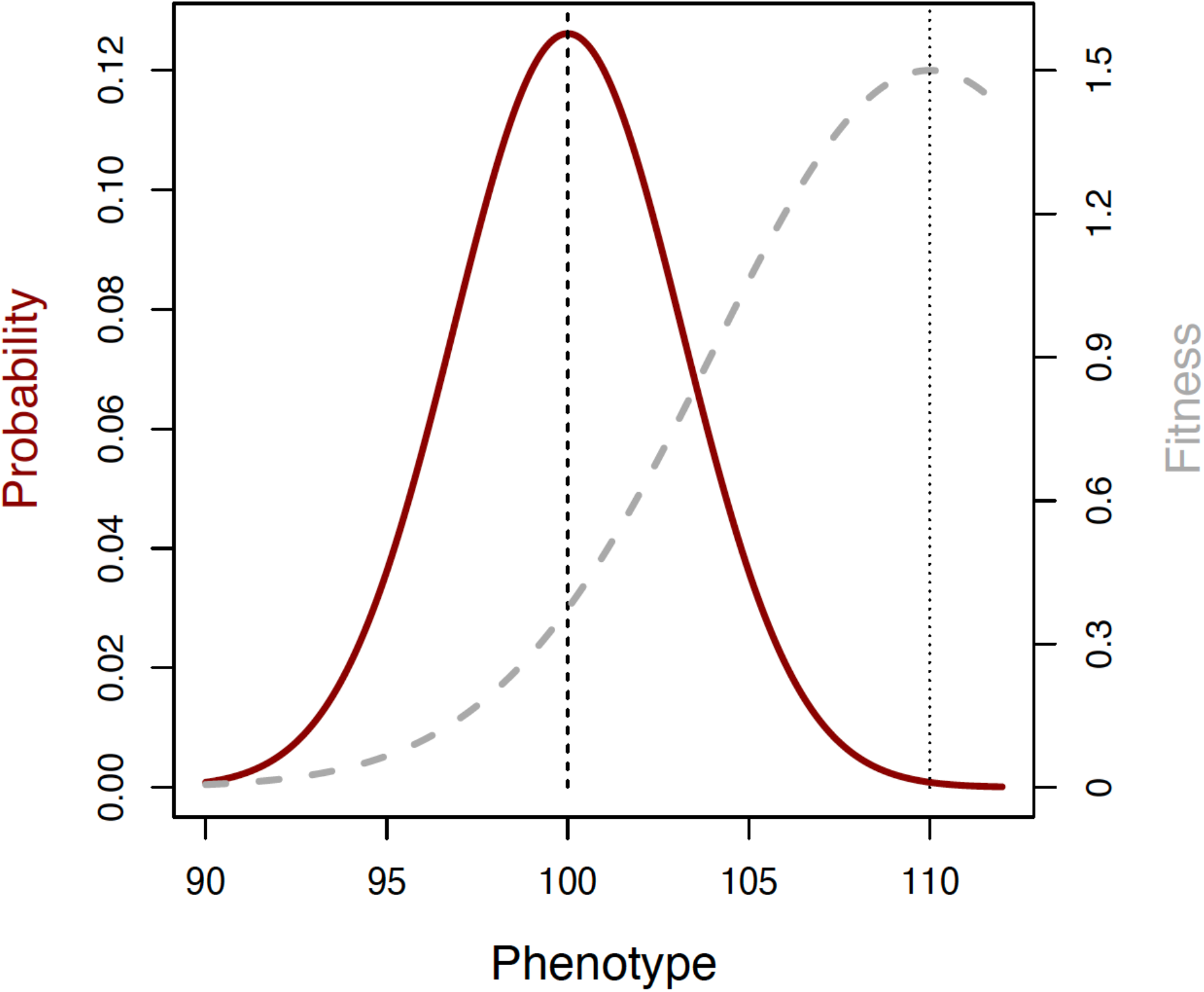
Fitness function and the phenotype distribution at the onset of selection in our determinstic model. The phenotype probability density distribuition is shown in red (left vertical axis). The vertical dashed line shows the initial mean phenothype 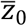. The gray dashed line represents the Gaussian fitness function with standard deviation *c* = 6 (right vertical axis). The vertical dotted line shows the new optimum phenotype *θ*. Integrating over the product of the phenotype and fitness functions [see eq. (6)] yields the mean intrinsic fitness 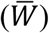 in the population (i.e., ignoring effects of population density).

The strong effect of the initial frequency of large-effect alleles on the temporal dynamics of *h*^2^ means it is crucial for the *p*_0_ values to be biologically meaningful. Large-effect alleles occur across a wide range of frequencies in natural populations (Barson et al. 2015; Johnston et al. 2013; Küpper et al. 2015; Thompson et al. 2019). For example, the *GREB1L* locus strongly affects seasonal timing of migration from the ocean to freshwater (ranging from Spring to Fall) in Chinook salmon and steelhead (Thompson et al. 2019). The allele associated with earlier entry into fresh water occurred at frequencies ranging from 0.002 to 0.488 across three populations (Thompson et al. 2019). Several mechanisms, including balancing selection (e.g., net heterozygous advantage or spatial variation in phenotypic optima), gene flow among populations with different phenotypic optima, and directional selection associated with historical environmental change can lead to large-effect polymorphisms occurring across a wide range of allele frequencies (Barson et al. 2015; Johnston et al. 2013). We therefore consider a broad range of initial frequencies of common beneficial alleles in our analysis of this model (*p*_0_ = 0.1, 0.25, 0.5, 0.75, or 0.9) at each of *n* loci that affect the selected trait. Evolutionary potential is determined by *p*_0_ in this scenario when 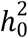 is held constant (Walsh and Lynch 2018). Varying *p*_0_ while holding 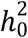 constant therefore allows us to evaluate the influence of evolutionary potential on population dynamics in this simplistic model. Note, however, that this analysis does not address historical factors that determine *p*_0_. Below we model initial allele frequencies as determined by historical mutation and selection in individual-based simulations.

While this model is useful for understanding population responses to selection, it makes some assumptions that are unlikely to hold in natural populations (e.g., no selection-induced linkage disequilibrium [LD]). LD among loci affecting a selected phenotype could be substantial, and may affect the pace of adaptation when multiple loci are involved and locus-specific selection is strong (Barton and Turelli 1991). This model (and previous similar models) also assumes that the selected phenotype is normally distributed. However, strong selection and/or large-effect loci might skew the phenotype distribution away from normality (Barton and Turelli 1991). We therefore repeated the above analyses, this time implementing an explicit simulation-based model of genotype and the phenotype distributions for this model (details in Supplementary Materials). The simulated phenotypes were approximately normally distributed (Figure S1).

### A stochastic, individual-based simulation model of population responses to environmental change

While deterministic models, such as the ones described above, are useful for understanding expected responses to selection, they do not incorporate the potentially important effects of evolutionary and demographic stochasticity on population responses to a changing environment. We therefore developed a stochastic, individual-based simulation model of evolution and population dynamics under environmentally-induced selection. This model simulates populations forward-in-time with density-dependent fitness and viability selection on a quantitative trait. The initial population size was set to *N*_0_ = 500 individuals, with a carrying capacity of *K* = 1,000 individuals. Fitness was density-dependent and followed the discrete logistic model of population growth in eq. (10) above. Mates were paired at random, with no self-fertilization allowed. The number of offspring per breeding pair was Poisson distributed (i.e., assuming randomly distributed fecundity among breeding pairs) with an arbitraily assigned mean and variance of 4 offspring. Alleles were transferred from parent to offspring following Mendelian probabilities.

#### Simulating the selected phenotype

The selected phenotype had an initial variance of *V*_z_ = 10 and an initial heritability of 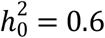. Individual *i*’s phenotype was

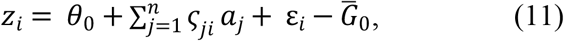

where *θ*_0_ is the specified optimum (equal to the initial mean) phenotype in the first generation, 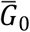 is the mean additive genetic value among individuals (i.e., the second term in [10]) in the first generation, 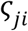 is individual *i*’s count of the allele conferring a larger phenotype at the *j*^th^ of *n* loci, and *aj* is the phenotypic effect of the positively selected allele (i.e., the allele conferring a larger phenotype) at the *j*^th^ locus, and the environmental effect *ε_i_* is drawn at random from a normal distribution with mean = 0 and variance = *V*_E_. As in eqs. (3) and (8), the 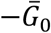 term in (11) ensures that the selected phenotype distribution is centered around *θ*_0_ in the first generation. We simulated phenotypes as a function of 1 or 2 large-effect loci or 100 small-effect loci, each with the same initial beneficial allele frequency *p*_0_, and effect size *a* (consistent with the deterministic models above). Each locus had additive phenotypic effects and there was no epistasis.

#### Fitness as a function of phenotype

Each population was subjected to viability selection on the simulated phenotype. The expected (deterministic) fitness (*w*) for each individual in generation *t* was calculated as in equation (1) above. The mean deterministic fitness in generation 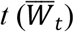, *N*_*t*_, and *K* were applied to equation (10) to find the deterministic expected population size in generation *t* + 1 (*N*_exp,t+1_, the total expected number of offspring left by generation *t*). The mean probability of surviving to breeding age among individuals in generation *t* was then calculated as

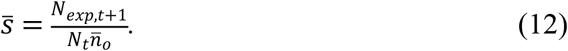

The number of individuals in generation *t* surviving to maturity was calculated as

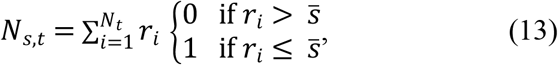

where *R*_*i*_ is a number selected at random from a uniform distribution bounded by 0 and 1 (using the *runif* function in R). This is equivalent to a random draw from binomial distribution with parameters *N*_*t*_ and 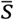. *N*_*s,t*_ individuals surviving to maturity in generation *t* are then selected at random from *N*_*t*_ individuals, with sampling weight *w,* such that individuals with *z* closer to *θ* are more likely to survive to maturity. We calculated the extinction rate each generation as the proportion of 500 replicate simulated populations with < 2 individuals remaining. A flow chart summarizing the structure of the individual-based simulation model is shown in Figure S2.

### Simulations of different life histories, heratibilities, and allele frequency distributions

The simulations above assumed that all of the positively-selected alleles conferring a larger phenotype have the same *p*_0_ and equal phenotypic effects. A more realistic situation is likely where a selected phenotype is governed by both large- and small-effect loci across a wide range of initial allele frequencies. We therefore modified our individual-based simulation model to evaluate the effects of genetic architecture on population responses to selection when both large- and small-effect loci with a wide range of initial allele frequencies were present. The selected phenotype had an initial heritability of 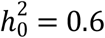.

We ran simulations with and without a single large-effect locus. For simulations with a large-effect locus, we attributed 90% of the *V*_G_ to the large-effect locus as 2*pqa*^2^ = 0.9*V*_G_. The remaining 10% of the *V*_G_ was split evenly among the other 99 loci. For simulations without a large-effect locus, the *V*_G_ was split evenly among all 100 loci. The residual phenotypic variance (*V*_E_) was attributed to random environmental differences among individuals (*V*_E_ = *V*_z_ − *V*_G_; *V*_G_ = *h*^2^*V*_z_). The initial frequency of the positively selected, large-effect alleles was drawn at random from a uniform distribution ranging from 0.05 to 0.95. We set these *p*_0_ limits to avoid extremely large phenotypic effects (i.e., extreme values of *a*) at large-effect loci, while incorporating a broad range of large-effect allele frequencies as observed in natural populations (Barson et al. 2015; Thompson et al. 2019). The *p*_0_ values at the small-effect loci were drawn at random from a beta distribution with parameters *α* and *β* each set to 0.5 which results in a typical U-shaped allele frequency distribution where most loci had the minor allele at low frequency (Kimura 1984) [p. 147].

We parameterized these simulations to approximately mimic two divergent life histories including high survival combined with low fedundity (e.g., large mammals; Mduma et al. 1999) and low survival combined with high fedundity (e.g., free-living corals; Fadlallah 1983) to determine if life history strategy affected the results. The maximum fitness (*W*_max_ = the expected reproductive success of a perfectly adapted individual at very low population density) was *W*_max_ = 1.5 for large mammals (mean number of offspring per breeding pair = 4, survival to maturity probability = 0.75), and *W*_max_ = 1.3 for corals (mean number of offspring per breeding pair = 26; survival to maturity probability = 0.1). Note that *W*_max_ is equivalent to the geometric population growth rate (*λ*) for a perfectly adapted population with *N* very near zero. We assumed *N*_0_ = 500 and *K* = 1,000, and *N*_0_ = 10,000 and *K* = 20,000 for simulations of approximate large mammal and coral life histories, respectively. We initially ran 1,000 coral and large mammal simulation repetitions (500 with a large-effect locus, and 500 with a polygenic trait architecture) to evaluate the effects of genetic architecture on the population responses to selection associated with the shifted phenotypic optimum. We ran 1,500 additional simulations with a large-effect locus and 99 small-effect loci affecting the selected phenotype to determine how *p*_0_ of a large-effect locus affected population dynamics.

We varied the parameter values of our individual-based simulations using a large mammal life history to test whether our findings hold across a range of other scenarios. For example, the size of the shift in *θ* (particularly with reference to the width of the fitness function [*c*]) is a key parameter as it determines the effect of an environmental change on fitness. Our main analyses considered an sudden increase in *θ* of 10 units (Figure 1), such that the new optimum phenotype was in the far right tail of the initial phenoytpe distribution and elicited a substantial decrease in fitness (see Results). We added a scenarios where *θ* shifted by only 5 and 7.5 units to test the effect of trait genetic architecture on population dynamics associated with weaker selection arising from smaller changes in the optimum phenotype. We also accounted for potential effects of variation in other parameters by simulating lower and higher initial heritability of the selected trait (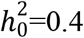 and 0.8), gene flow from a population where *θ* was held constant at *θ*_0_, weaker effects at large-effect loci (i.e., the large-effect locus being responsible for 50% and 70% of *V*_G_), a stochastic linear temporal increase in *θ* (Lynch and Lande 1993)(instead of a sudden shift as above), selection with linked loci (on 10 chromosome pairs), and plasticity in the selected phenotype. The methodological details of these simulation scenarios are described in the Supplementary Materials.

Lastly, we developed another individual-based simulation model that explicitly accounts for effects of historical factors on the allele frequency distribution at the onset of environmental change and subsequent responses to selection. These simulations used a long burnin period (≥ 1,000 generations) to allow the *V*_G_ to reach approximate mutation-drift-selection equilibrium before shifting the phenotypic optimum. The details of this model are described in the Supplementary Materials.

### Effects of the short term selection limit on population dynamics

Effects of genetic architecture on responses to environmental change may be driven largely by variation in the potential for populations to evolve rapidly. To test this, we defined a short term selection limit (*L*) and quantified its relationship to population viability in our individual-based simulations with different genetic architectures underlying the selected phenotype. We defined *L* as the expected adaptive change in the mean phenotypic 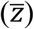 over *t* generations, assuming that the difference between 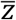 and the phenotypic optimum (*θ*) is constant through time (i.e., an increase in 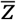 over a generation results an equivalent increase in *θ*), and that natural selection is the only driver of phenotypic evolution. *L* therefore measures the potential of a population to evolve in response to selection over the short term under the idealistic conditions of consistently strong selection and no genetic drift.

We used numerical methods to calculate the initial *L* at the beginning of each simulation repetition (*L*_0_) in the scenarios with a large mammal life history and no mutation. We first used equations 4–7 to calculate the expected change in the allele frequencies and mean phenotype over the first *t* = 10 generations under the conditions outlined above. We chose *t* = 10 generations because most simulated populations that persisted began increasing in size by the tenth generation (see Results). *L*_0_ was then calculated as the difference between the predicted mean phenotype at *t* = 10 generations and the mean phenotype at the beginning of the simulation.

We used regression analysis to measure the relationship between *L*_0_ and population viability. The *glm* function with a logit link function in R was used to fit generlized linear models (GLMs) with population persistence as the reponse (coded as 0 for extinct populations, and 1 for populations that persisted for 80 generations) and *L*_0_ was the predictor. We fitted separate GLMs for the 500 simulations with a large-effect locus, and for the 500 simulations with a polygenic selected trait in each simulation scenario. The odds ratios from the GLMs were used to measure the size of the effect of *L*_0_ on population viability. We also analyzed the data from all simulation scenarios (both with and without large-effect loci) combined in a single GLM to evaluate the influence of *L*_0_ on population persistence across all of the analyzed scenarios and genetic architectures. Finally, we quantified the temporal dynamics in *L* across the first 30 generations in each simulation scenario to determine how the potential for rapid evolution changed throughout the selection response under different genetic architectures.

### Statistical analysis of extinction rate

We constructed 95% percentile bootstrap confidence intervals (Efron and Tibshirani 1994) for the proportion of extinct populations in each simulated scenario. First, we randomly resampled *η* simulated data sets 1,000 times, with replacement, from the *η* original simulation repetitions. For each of the 1,000 bootstrap samples, we calculated the proportion of the *η* resampled populations that were extinct (*N* < 2 individuals) in each of the 80 generations. We constructed the 95% bootstrap confidence intervals for the extinction rate for each of the 80 generations as the 2.5% and 97.5% quantiles from the bootstrap distributions.

## RESULTS

### Deterministic predictions of evolutionary and demographic responses to environmental change

Results from our deterministic model suggest that the genetic architecture underlying the *h*^2^ of a selected trait strongly affects the evolutionary and demographic responses to a sudden environmentally-induced shift in the phenotypic optimum *θ*. First, phenotypic evolution and population growth after the onset of selection were highly dependent on the initial frequency *p*_0_ of large-effect alleles, but relatively insensitive to *p*_0_ when many small-effect loci were involved (Figure 2). Populations with already-frequent large-effect benficial alleles did not have enough evolutionary potential to remain viable. For example, populations with a single large-effect locus and *p*_0_ ≥ 0.5 all went extinct before 30 generations as they were unable to approach the new phenotypic optimum. However, population size *N* eventually approached carrying capacity *K* in populations with *p*_0_ < 0.5. The time to reach *N* ≈ *K* was approximately 25 generations longer when there was a single large-effect locus and *p*_0_ = 0.25 compared to *p*_0_ = 0.1 (Figure 2A). With 2 large-effect loci, the expected time to reach *N* ≈ *K* was nearly identical for *p*_0_ = 0.1 and *p*_0_ =0.25. Populations with 2 large-effect loci recovered slowly with *p*_0_ = 0.5, and went extinct by 20 generations with *p*_0_ > 0.5 (Figure 2B).

**Figure 2.**
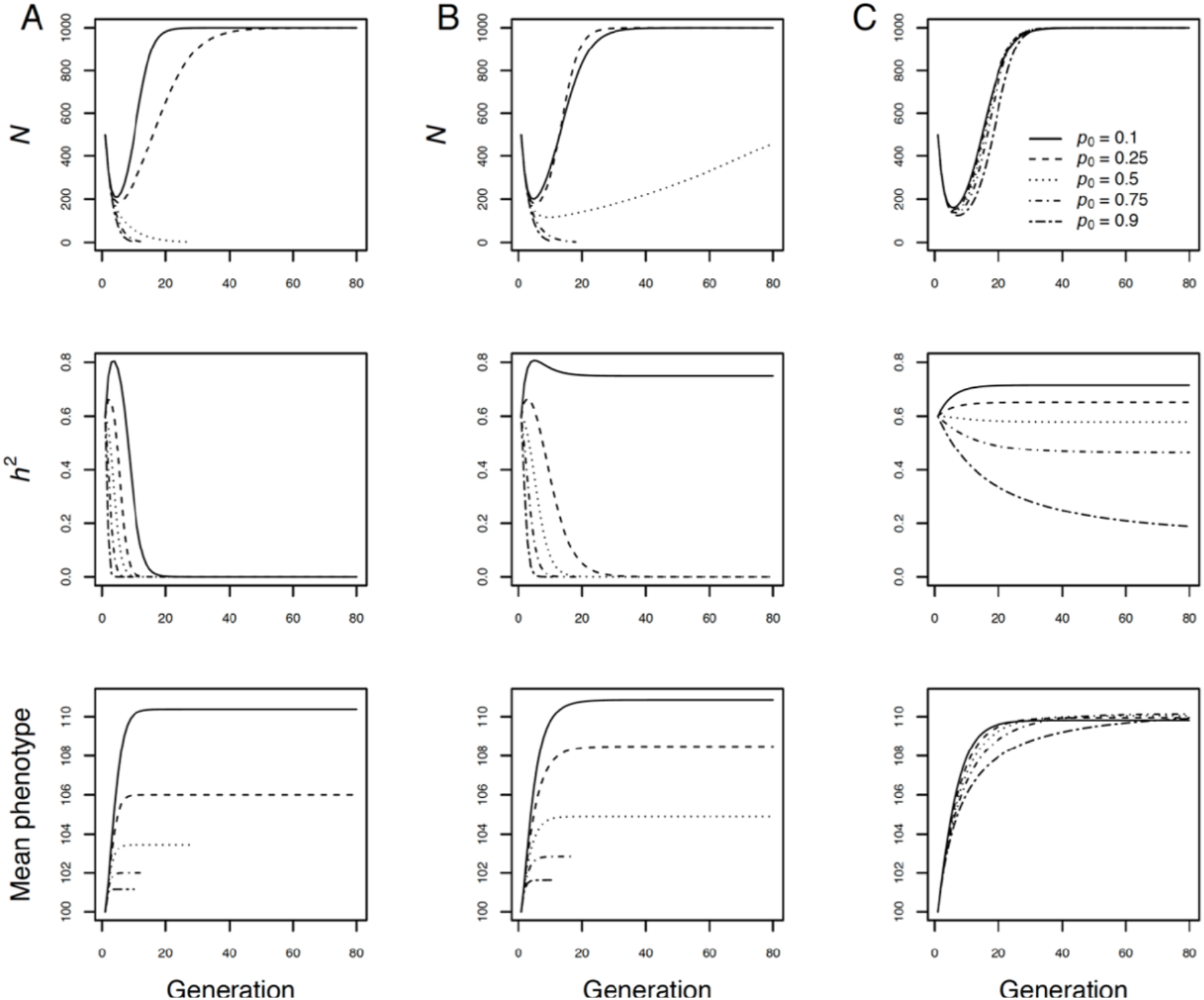
Deterministic predictions of population size (*N*, top row), trait heritability (*h*^2^, middle row), and mean phenotype (bottom row) through time in density-regulated populations with a single large-effect locus (**A**), two large-effect loci (**B**), and 100 small-effect loci (**C**) affecting a quantitative trait under selection after a sudden environmental change. Initial population size was *N* = 500 with carrying capacity of *K* = 1,000, and the initial heritability was 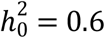 in all cases. The optimum phenotype shifted from *θ*_0_ = 100 to *θ*_1_ = 110 in the first generation. Line types indicate the initial frequencies of the positively selected allele(s) conferring a larger phenotype, as indicated in the legend.

The rate of adaptation and recovery of population size was much less affected by *p*_0_ when the selected trait was polygenic, with the phenotype approaching the new phenotypic optimum *θ*_1_, and *N* approaching *K* for all values of *p*_0_ (Figure 2C). Results from analyses of this model with initial heritability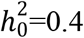 and 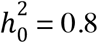 were qualitatively equivalent to the results presented here (Figures S3, S4). Repeating these analyses with simulated phenotypes to account for selection-induced LD and any deviation from the assumed normal phenotype distribution did not substantively affect the results (Supplementary Materials, Figure S5).

### Stochastic, individual-based simulations of evolutionary and demographic responses environmental change

Similar to the deterministic results, our individual-based simulations show that the lowest initial positively-selected, large-effect allele frequency (*p*_0_ = 0.1) conferred substantially increased adaptation, demographic recovery, and a lower extinction rate compared to beneficial, large-effect alleles with higher *p*_0_ (Figure 3A, 3B). The phenotypic response to selection was larger over the long run for the polygenic architecture than with large effec loci for all *p*_0_. This led to the polygenic architecture conferring lower extinction rate and larger *N* on average compared to the large-effect genetic architectures for all *p*_0_ values except *p*_0_ = 0.1, in which case the large-effect loci resulted in faster adaptive phenotypic evolution and population size recovery from selection (along with lower extinction rates) compared to the polygenic architecture. Repeating these individual-based simulations with 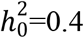 and 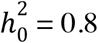 generated results that agreed qualitatively with those presented here 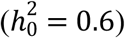 (Figures S6-S9).

**Figure 3.**
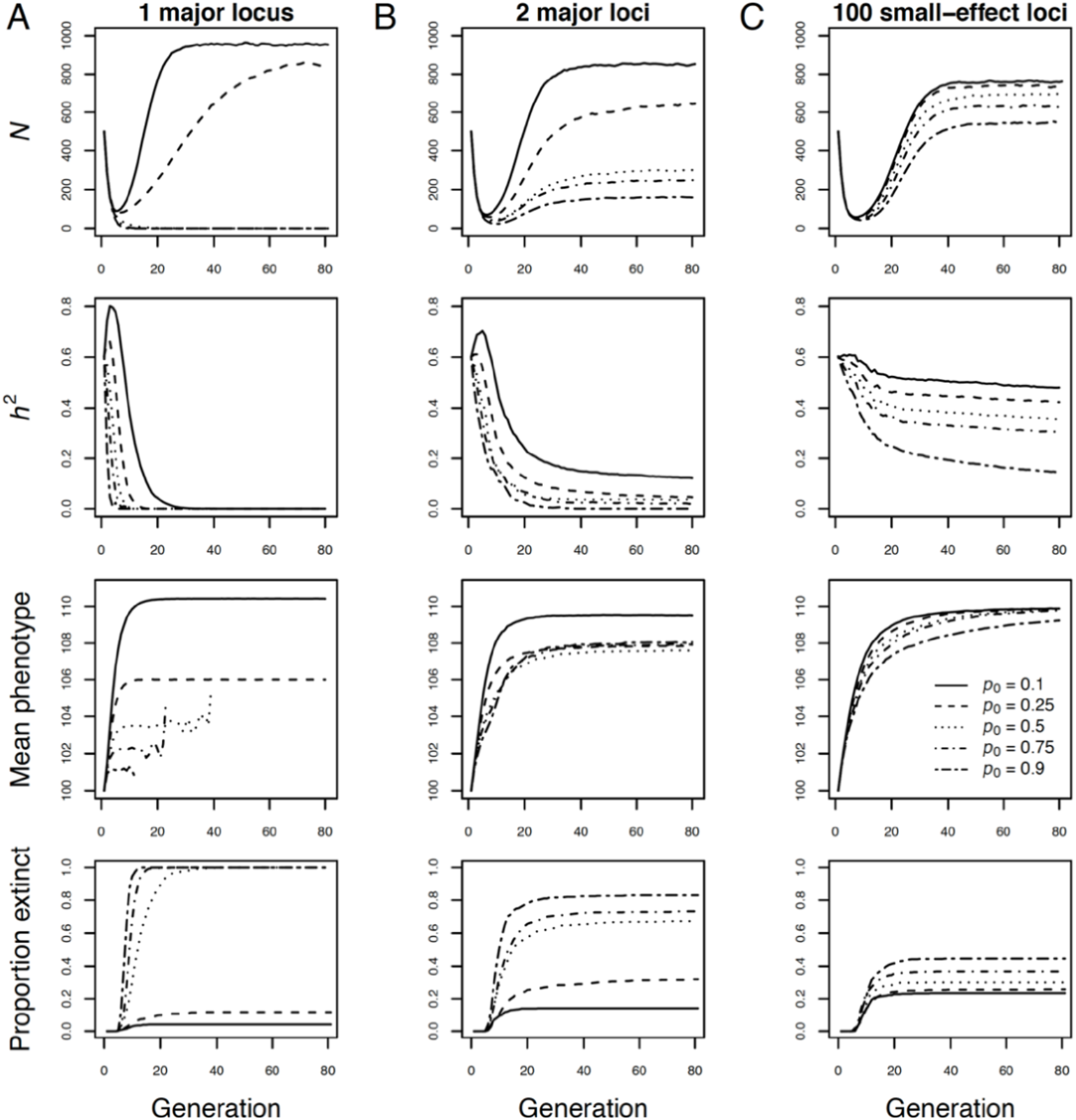
Individual-based simulations of evolutionary and population dynamics in density-regulated populations with a single large-effect locus (**A**), two large-effect loci (**B**), and 100 small-effect loci (**C**) affecting a quantitative trait under selection after a sudden environmental change. The optimum phenotype shifted from *θ*_0_ = 100 to *θ*_1_ = 110 in the first generation. Initial population size was *N*_0_ = 500, and capacity was *K* = 1,000. The initial heritability was 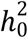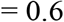 in all cases. Line types indicate the initial frequency of the positively selected allele(s) conferring a larger phenotype as indicated in the legend.

### Simulations of different life histories and allele frequency distributions

Results from our simulations of populations with variable *p*_0_ are similar to the simpler models presented above, with the very large-effect alleles conferring less adaption, smaller population sizes, and a higher extinction rate on average than when the selected trait was polygenic (Figure 4). Many populations with an already-frequent large-effect allele were unable to reach the new phenotypic optimum. Note though that some populations where the large-effect beneficial allele was initially rare overshot the phenotypic optimum. Populations with a polygenic selected trait more closely matched the new phenotypic optimum on average compared to the populations with a large-effect locus (Figure 4). The extinction rate at generation 80 was 2.0 times higher with the large-effect locus (64% extinction rate) compared to the polygenic architecture (32% extinction rate) in simulations assuming a large mammal life history. Similarly, the extinction rate was 2.7 times higher among populations with a large-effect locus (72% extinction rate) compared to the polygenic architecture (27% extinction rate) in simulations assuming a free living coral-like life history.

**Figure 4.**
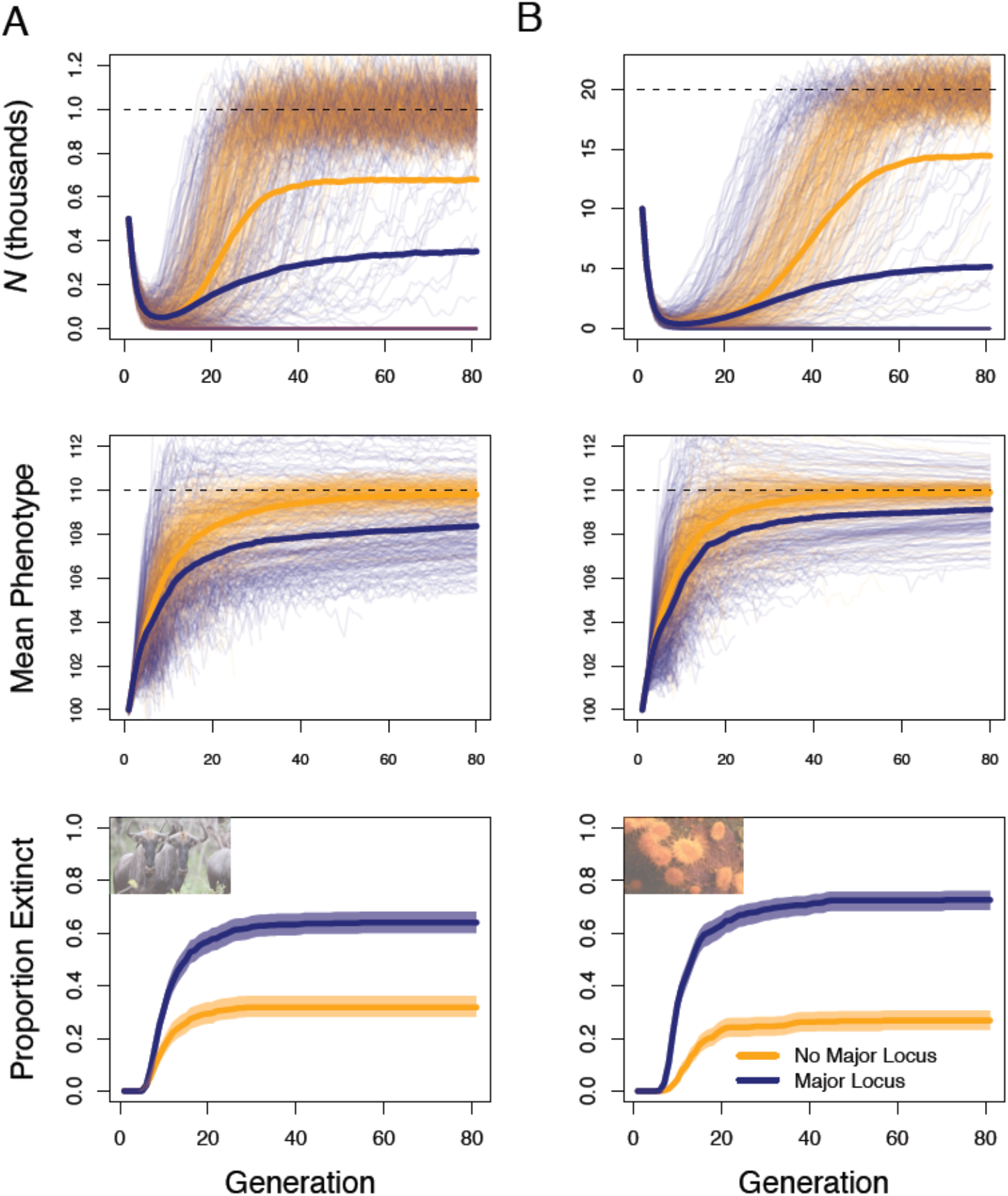
Effects of genetic architecture on phenotypic evolution and population dynamics in closed populations with life histories approximating large mammals (**A**), and corals (**B**). Results are shown in blue for populations with a large-effect locus, and in orange for populations where the selected trait was polygenic. The phenotypic optimum permanently shifted from its initial value *θ*_0_ = 100 to *θ*_1_ = 110 in generation one. The initial heritability was 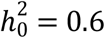. Thin colored lines show the population size (top row) and mean phenotype (middle row) through time. Thick colored lines show the mean population size and phenotype across all 500 repetitions. The bottom panels show the proportion of extinct populations through time, with percentile bootstrap 95% confidence intervals. The dashed black lines represent the carrying capacity in the top two panels, and *θ*_1_ in the middle panels.

These simulation results further suggest that *p*_0_ at large-effect loci strongly affects population dynamics (Figure 5). The average final population sizes were highest for both life histories when *p*_0_ was ~0.1-0.2. The lower average population growth with *p*_0_ < 0.1 is likely caused by rare, positively-selected alleles frequently being lost to genetic drift as the populations initially declined rapidly due to selection. The weaker evolutionary and demographic response in populations with already-frequent, large-effect beneficial alleles (Figure 4) resulted in lower population growth rates and eventual extinction in a large fraction of populations with *p*_0_ > 0.2. Strikingly, all of the populations with a coral life history and *p*_0_ > 0.5 went extinct by generation 80.

**Figure 5.**
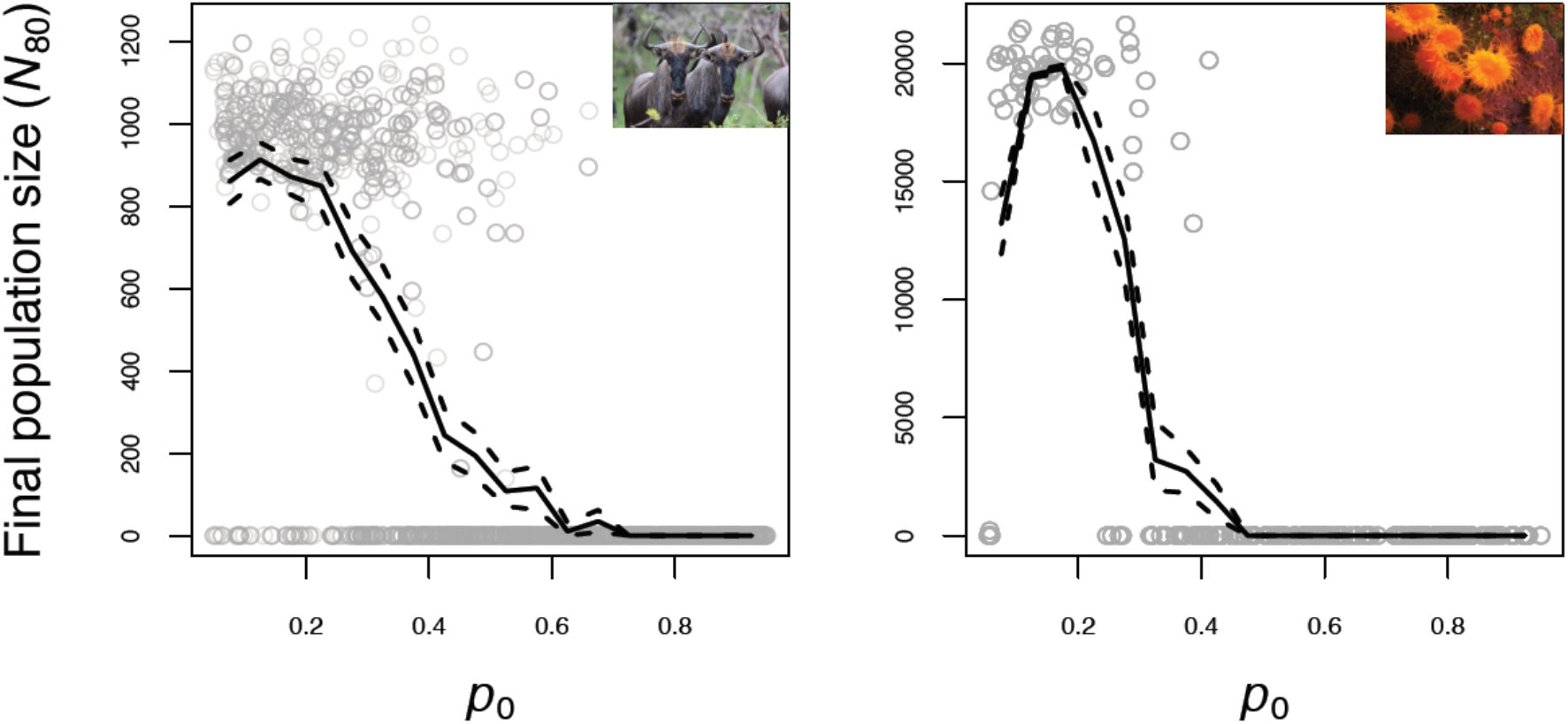
Effects of the initial large-effect allele frequency (*p*_0_) on final population size in simulations with approximate large mammal (**A**) and coral (**B**) life histories. The y-axis represents the final population size at generation 80 (*N*_80_), and the x-axis shows the large-effect allele *p*_0_. The solid lines represent the mean *N*_80_ across 2,000 simulation repetitions in non-overlapping *p*_0_ windows of width 0.05. Dashed lines are 95% percentile bootstrap confidence intervals for mean *N*_80_.

Polygenic architectures conferred higher population viability on average compared to when large-effect loci were present for all of the alternative simulation scenarios: lower and higher initial heritability than above, gene flow from a population with a stationary phenotypic optimum, linked loci, weaker effect sizes at large-effect loci, a stochastic linear temporal increase in *θ*, phenotypic plasticity, a smaller shift in *θ* (from 100 to 107.5), and with mutation and historical stabilizing selection (Supplementary Materials; Figures S6-S17). However, the increased evolutionary and demographic responses to selection associated with polygenic architecture was smaller when there was immigration from a population with a stationary *θ* (Figures S10 & S11), when the large-effect loci contributed a smaller fraction of the genetic variance *V*_G_ (Figure S12), and when the environmentally-induced shift in the optimum phenotype was smaller (see detailed results in Supplementary Materials). For example, the extinction rate at 80 generations was only 1.53 times higher with a large effect locus than for the polygenic architecture (compared to a 2-fold difference in the simulations of closed populations above) when there were 4 immigrants per generation from a population with a stationary phenotypic optimum (Supplementary Materials, Figure S9; similar results for 8 immigrants/generation are shown in Figure S10). The extinction rate for populations with a large-effect locus contributing only 50% of the *V*_G_ was only 1.28 times higher than in populations with a polygenic architecture (Supplementary Materials, Figure S12). For populations with a smaller shift in the phenotypic optimum (*θ*_0_ = 100 and *θ*_1_ = 107.5), the extinction rate was 8.6% in the populations with a large effect locus explaining 90% of the *V*_G_, and only 0.5% among populations with a polygenic selected phenotype (Figure S16). Less than 1% of all populations went extinct when the phenotypic optimum shifted from *θ*_0_ = 100 to *θ*_1_ = 105 (Figure S16).

### Effects of the short term selection limit on population dynamics

Population viability (i.e., persistence versus extinction) was statistically significantly associated with the initial short term selection limit *L*_0_ (*P* < 0.05) in seven out of the eight simulation scenarios with a large effect locus (Figures 6, S19-S26, Table S1). The only scenario with a large-effect locus where population viability was not statistically significantly associated with *L*_0_ was when the selected phenotype was strongly plastic (plasticity parameter *m* = 0.4, Figure S26) where the extinction rate was only 3%. The odds ratios from the GLMs of population persistence versus *L*_0_ in scenarios with a large effect locus ranged from 1.19 when there was a sudden shift in *θ*, no plasticity, and a major locus responsible for 50% of *V*_G_, to 2.94 when there was a sudden shift in *θ*, no plasticity, and a major locus responsible for 90% of *V*_G_ (Table S1). This translates to a 19% to 294% increase in the odds of population persistance per unit increase in *L*_0_.

**Figure 6.**
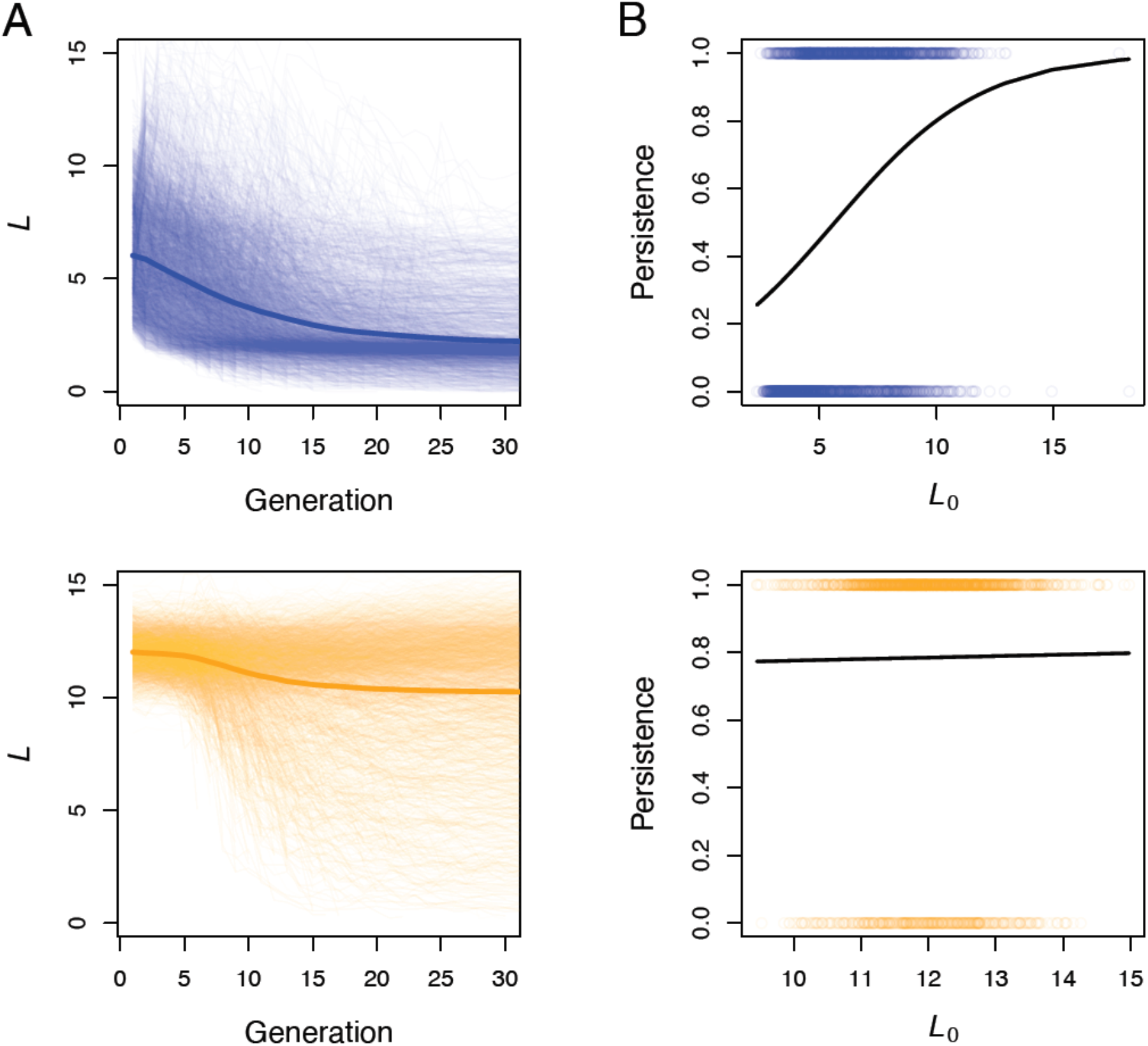
Temporal dynamics of the sort term selection limit (*L*) (**A**) and the relationship between population persistence (0 = extinct; 1 = population persisted) and the initial short term selection limit (*L*_0_) (**B**). The data are shown from all simulation scenarios where there was a large-effect locus (blue) and where the selected trait was polygenic (orange) combined. The thick colored lines in **A** represent the mean across all 4,000 individual simulation repetitions. The black lines in **B** are fitted logistic regression lines.

Population persistence was not statistically significantly associated with *L*_0_ (*P* < 0.05) in any scenario where the selected phenotype was polygenic (Table S2). The odds ratios from GLMs from simulations with a polygenic trait were centered around one, ranging from 0.85 to 1.25 (Table S2). The only scenario with a large-effect locus that had an odds ratio similar to its polygenic counterpart was when the the large-effect locus was responsible for 50% of the *V*_G_ (Figure S27, Tables S1 & S2).

Consistent with results from individual simualtion scenarios, population viability was not statistically significantly associated with *L*_0_ when analyzing simulations from all scenarios with a polygenic selected trait combined (*P* = 0.65, odds ratio = 1.03). However, population persistence was statistically significantly associated with *L*_0_ in our analysis of all simualtion scenarios combined with a large-effect locus combined (*P* < 2×10^−16^, odds ratio = 1.38). The GLM of population viability versus *L*_0_ across all simulated scenarios (i.e., all simulations with polygenic and major locus trait architectures combined) was statistically significant (*P* < 2×10^−16^) with an odds ratio of 1.25, meaning that a one unit increase in *L*_0_ was associated with a 25% increase in the odds of population persistance (Figure S27).

Polygenic trait architectures conferred larger short term selection limit *L* than genetic architectures with large-effect loci, both at the onset of selection and subsequently through the first 30 generations (Figures S19-S26; Figure 6). The average *L*_0_ across 500 simulation replicates was approximately 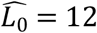 in all simulation scenarios with a polygenic selected trait. 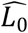 ranged from 5.21 to 5.36 among simulations with a large-effect locus responsible for 90% of the *V*_G_. However, the 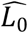 was condiserably higher for simulations where the large-effect locus was responsible for 70% of the 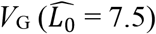, and 50% of the 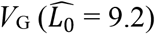 (Figures S20 & S21).

## Discussion

The results from the wide range of analyses above suggest that the genetic architecture underlying the *h*^2^ of a selected trait can strongly affect population viability during environmental change. Understanding of the effects of environmental change on population viability will be advanced by accounting for the strong effects of trait genetic architecture on evolutionary and population dynamics. Polygenic architectures on average conferred higher evolutionary potential, more consistent evolutionary responses to selection, and increased population viability compared to when the selected trait was governed by large-effect loci. When loci with large phenotypic effects are present, the initial frequency of large-effect beneficial alleles can strongly affect population responses to selection. Large-effect loci appear to confer adaptation and demographic recovery that is similar or higher than with polygenic architectures only when the positively selected alleles are initially infrequent (Figure 4). Additionally, while predicting how wild populations will respond to ongoing rapid environmental change remains challenging, the models and results presented here can inform future theoretical and empirical efforts to understand eco-evolutionary dynamics and the extent of the ongoing extinction crisis.

The influence of genetic architecture on variation in population responses to environmental change will depend on how often fitness traits have loci with large enough effects to alter *h*^2^ during bouts of adaptation. Recent results from several taxa, including mammals (Barson et al. 2015; Epstein et al. 2016; Jones et al. 2018; Kardos et al. 2015), salmonids (Barson et al. 2015; Pearse et al. 2019; Thompson et al. 2019), and birds (Lamichhaney et al. 2015; Lamichhaney et al. 2016) suggest that very large-effect alleles often influence fitness-related traits in wild populations. Interestingly, variation in seemingly complex fitness-related traits that are often assumed to be polygenic, such as horn size (a sexually-selected, condition-dependent trait) (Johnston et al. 2013), migration timing (Thompson et al. 2019), propensity to migrate (Pearse et al. 2019), and age at maturity (Barson et al. 2015), has in some cases turned out to be driven almost entirely by variation at large-effect loci. It is therefore crucial to quantify the effect sizes and allele frequencies at loci with large effects when they are present in systems where future eco-evolutionary dynamics are of interest (Funk et al. 2018; Yang et al. 2014).

It can be difficult to predict or measure the frequency of alleles with large beneficial effects under rapid environmental change. For example, large-effect alleles for traits subjected to historical balancing selection, are likely to be at intermediate frequencies (Llaurens et al. 2017). Recent large-effect mutations are likely to be found at low frequencies. Previously neutral or nearly-neutral alleles that affect fitness in new conditions are likely to be found across the entire spectrum of allele frequencies. Fortunately, increasingly efficient DNA sequencing and improving approaches for conducting genotype-phenotype association analysis provide the tools necessary to estimate *h*^2^, and to identify large-effect loci (and to estimate their allele frequencies) where they exist.

Why do polygenic architectures usually confer increased population viability compared to genetic architectures including large-effect loci? This pattern arises in part from a slower and less variable decline in *h*^2^ during adaptation for polygenic traits than for traits with large-effect loci (Figures S3-S6, S8). The rapid decline in *h*^2^ when beneficial alleles with large effects are already common, and the frequent loss of initially rare large-effect alleles means that there is a narrow window of *p*_0_ where traits with large-effect architectures are likely to evolve in response to selection as fast or faster than polygenic traits. Holding the initial heritability constant, the potential for adaptive phenotypic change is considerably smaller when large-effect loci are present compared to a polygenic architecture (Walsh and Lynch 2018)(Figure 6). It appears that large effect loci often do not confer enough adaptive potential over the short term to accomodate large, rapid shifts in phenotypic optima. Additionally, evolutionary and demographic responses to selection appear to be more stochastic in populations with large-effect loci (Figure 4). This suggests that reliably predicting population responses to selection will be more difficult when large-effect loci are present, particularly when the initial large-effect allele frequency is not known precisely. These results highlight the importance of identifying large-effect loci where they exist, and using information on their effect-sizes and allele frequencies along with 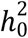 in models predicting demographic responses to environmental change. Predictions of population responses to selection are likely to be misleading if they do not account for the strong effects of genetic architecture on the temporal dynamics of *h*^2^ and adaptation.

Understanding how populations will respond to environmental change is both challenging and vitally important in conservation and evolutionary ecology (Urban et al. 2016). Reliable predictions of how biodiversity will respond to large-scale environmental change are necessary to efficiently prioritize scarce conservation resources and to develop effective conservation strategies. Improved understanding of vulnerability to environmental change could also advance strategies to conserve vital natural and agricultural resources (Aitken and Whitlock 2013; Flanagan et al. 2018; Funk et al. 2018), for example by identifying populations and species to prioritize for conservation action. However, there are substantial obstacles to reliably predicting responses to selection. The complex and interacting effects of environmental stochasticity, genotype-by-environment interactions, phenotypic plasticity, pleiotropy, dominance interactions, gene flow, simultaneous selection on correlated traits, and changing community structure (i.e., species interactions) can all strongly affect adaptation and population dynamics, but are also difficult to measure and to forecast into the future. Consequently, reliable predictions of population responses to environmental change in the wild will be difficult to achieve, even in well-studied systems where the heritability, genetic architecture, and fitness effects of the relevant phenotype(s) are known. We therefore encourage caution when attempting to predict eco-evolutionary dynamics under climate change and other human-driven environmental changes.

While recognizing the difficulties involved, our results suggest that integrating genomic, classical quantitative genetic, and population viability analyses (e.g., applying the modelling approaches used here) is likely the most promising way forward to increased understanding the impacts of human-driven environmental change on population dynamics and extinction. Predictions of evolutionary and demographic responses to selection based only on trait loci detetected with genomic analyses will often be unreliable because a substantial fraction of phenotypic variation will frequently be explained by many undetected loci with small effects (Shaw 2019). We also argue that predictions based solely on classical quantatitive genetics approaches (Shaw 2019) will also frequently perform poorly because the selection response with large effect loci deviates strongly from expectations arising from the infinitesimal model of inheritance. Integrating genomic information (i.e., the genetic basis of phenotypic variation) into quantitiative genetic and population viability analyses will almost certainly improve predictions of responses to selection. Incorporating such ‘genomically-informed’ quantitative genetic approaches into population projection models has the potential to improve understanding of the impact of environmental change on population dynamics and extinction.

## Acknowledgements

M.K. was supported by Montana Fish, Wildlife & Parks. G.L. and M.K. were supported partially by National Science Foundation grant DoB-1639014, and National Aeronautics and Space Administration grant NNX14AB84G. We thank Fred W. Allendorf, Matthew Jones, L. Scott Mills, Alden Wright, Jeff Hard, Robin Waples, Michael Morrissey, and anonymous reviewers for helpful discussions and/or comments on previous versions of the manuscript.

## Data Availability

This manuscript is based on mathematical models and simulations and does not include any empirical data. R packages implementing the deterministic and stochastic models, and R scripts that replicate the simulations, statistical analyses, and figures are freely available at https://github.com/martykardos/AmNat_geneticArchitecture.

## Author Contributions

M.K. conceived the study. M.K. and G.L. designed the study. M.K. wrote and analyzed the mathematical models and simulations. M.K. and G.L. wrote the paper.

