## Supplementary Materials for "The genetic architecture of fitness drives population viability during rapid environmental change"

Marty Kardos*
Flathead Lake Biological Station
Division of Biological Sciences
University of Montana

Northwest Fisheries Science Center
National Marine Fisheries Service
National Oceanic and Atmospheric Administration,
Seattle, WA, USA


Gordon Luikart
Flathead Lake Biological Station
Division of Biological Sciences
University of Montana


**Effects of gene flow on the influence of genetic architecture on population viability**

Previous work suggests that selection with gene flow can favor the evolution of genetic architectures that have large-effect loci[1]. The presence of large-effect loci may therefore confer more robust evolutionary and demographic responses to selection when there is gene flow from populations with a different phenotypic optimum. We ran simulations equivalent to our individual-based simulations with a large mammal-like life history in the main text, except with immigration (either 4 or 8 immigrants / generation) from a population where the phenotypic optimum remained at 100 instead of shifting to 110 in the first generation.

The polygenic trait architecture conferred higher population viability in populations with immigration from a population with a different phenotypic optimum. The extinction rate was 55% among populations with a polygenic architecture, and 76% in populations with a large-effect locus when the immigration rate was four individuals per generation (Figure S10). With a higher immigration rate (8 individuals per generation), the extinction rate was 48% among populations with the polygenic architecture, and 74% in populations with a large-effect locus (Figure S11).

**Effects of major locus effect sizes on population viability**

We then evaluated the influence of the major locus effect sizes on evolutionary and demographic responses to selection. We ran our simulations of large mammals with the large-effect locus responsible for 50%, 70% and 90% of $h_0^2$ and compared the results to simulations where the selected trait was polygenic. In all cases, the responses to selection were stronger for the polygenic architecture than when there was a large-effect locus. The extinction rates were 1.28, 1.84, and 2.01 times higher with large-effect loci responsible for 50%, 70%, and 90% of $h_0^2$, respectively, than for the polygenic architecture (extinction rate = 0.32, Figure S12).

**Effects of genetic architecture on population viability with a temporally increasing and stochastic phenotypic optimum value**

Our analyses in the main text of the paper focused on the simple scenario where selection is induced by a sudden and permanent environmental change. However, some environmental change is often stochastic and spread out over long periods of time. We therefore used our individual-based simulation model with a large mammal-like life history to evaluate the effects of genetic architecture on population dynamics under this type of environmental change. We ran simulations equivalent to those shown in Figure 4A in the main text, except here the phenotypic optimum increased linearly with time (from 100 to 110) over the first *x* generations, with an error term to incorporate temporal stochasticity. The phenotypic optimum value for generation *t* was calculated as

$$\theta_t = \begin{cases} \theta_0 + \frac{t}{x}(\theta_{max} - \theta_0) + \varepsilon & \text{if } t < x \\ \theta_{max} + \varepsilon & \text{if } t \geq x \end{cases} \tag{S1}$$

where $\theta_0$ is the initial phenotypic optimum, $\theta_{max}$ is the final deterministic phenotypic optimum value, and $\varepsilon$ is normally distributed with mean of zero and standard deviation of 2. We ran 500 simulation replicates with *x* = 10 generations, and also with *x* = 20 generations. The results are shown below in Figure S13, and were qualitatively equivalent to those presented in Figure 4A in the main text.

**Effects of genetic architecture on population viability with linked loci**

Our analyses in the main text assumed that loci were unlinked. Linkage disequilibrium arising from having linked fitness-related loci can affect the response to selection[2]. We therefore ran simulations equivalent to those shown in Figure 4A, except here we placed the selected loci randomly across 10 chromosomes, each with a genetic length of 50 cM and physical length of 100 Mb, randomly selected crossover locations, and a Poisson distributed number of crossovers per meiosis. The results are shown in Figure S14 below, and were nearly identical to those in Figure 4A.

**Effects of the genetic architecture on population viability with phenotypic plasticity**

A previous theoretical population genetic model suggested that loci with large effects may provide higher adaptive potential and population viability when the selected phenotype is plastic[3]. We therefore tested effects of genetic architecture in the presence of phenotypic plasticity. We used a model of plasticity similar to that of Nunney[3]. We modified equation (11) from the main text to incorporate a linear effect of the optimal phenotype on the realized phenotype as

$$z_i = \theta_0 + \sum_{j=1}^{n} c_{ji}\, a_j + m\left(\theta_t - \left[\theta_0 - \bar{G}_0 + \sum_{j=1}^{n} c_{ji}\, a_j\right]\right) - \bar{G}_0 + \varepsilon_i, \qquad \text{(S2)}$$

where $m$ is the rate change in expected phenotype per unit difference between the optimal phenotype and individual $i$'s expected phenotype in the absence of plasticity. The plasticity model is shown with examples in Figure S18 below. We ran 500 simulation replicates equivalent to those described above (where there is a stochastic and linear increase in $\theta$ with time [$x = 10$]) except here we considered values of $m = 0.1$, 0.2, and 0.4. The results from these simulations are consistent with the results above and in the main text (e.g., Figure 4). The polygenic architecture on average conferred higher population sizes and a lower extinction rate than when there was a large-effect locus in simulations with $m = 0.1$, and $m = 0.2$ (Figure S15). The extinction rate was <0.05 for both genetic architectures when phenotypic plasticity was strong ($m = 0.4$).

**Simulations with mutation and historical stabilizing selection**

The individual-based simulations above made simplifying assumptions regarding the allele frequency distribution at the onset of environmentally-induced selection. To account for effects of historical selection on the allele frequency distribution, we ran simulations with a burnin period where alleles affecting the selected trait arose by mutation, and then evolved by genetic drift and stabilizing selection on the quantitative trait prior to an environmentally-induced change in $\theta$ that challenged population persistence. These simulations assumed the large mammal life history described in the main text, and began with constant population size of $N_0 = 500$, phenotypic variance due only to environmental effects ($V_e = 4$), and no alleles segregating at loci that affect the trait. We used the stochastic house-of-cards model (Bürger and Lande 1994; Turelli 1984) to parameterize the mutation rate giving rise to the polygenic part of the $V_G$ ($V_{G,poly}$). We drew the effect sizes of new mutations from a flat distribution [following Bürger & Lande (1994)] ranging from $\alpha = -0.5$ to $\alpha = 0.5$. The necessary mutation rate was calculated as

$$\mu = \frac{V_{G,poly}\left(1+\sigma_\alpha^2 \frac{N_0}{c^2+V_e}\right)}{4\mathbb{N}\sigma_\alpha^2 N_0}, \qquad \text{(S3)}$$

where $\sigma_\alpha^2$ is the variance of the flat distribution from which $\alpha$ values were drawn, $\mathbb{N} = 1 \times 10^6$ is the number of unlinked nucleotides where mutations can occur (Bürger and Lande 1994)and $c = 6$ as in the analyses above.

We ran 500 simulations with large-effect mutations as follows. The polygenic mutation rate was set to $\mu = 9.35 \times 10^{-9}$ after setting $V_{G,poly}$ in (13) to 0.6 (10% off the eventual $V_G$). A single large effect mutation with effect size $\alpha$ was drawn randomly from a flat distribution ranging from $\alpha = 3$ to $\alpha = 8$ was introduced in every generation when there was no large effect allele already in the population. This range of $\alpha$ values was chosen as populations with $\alpha > 8$ almost always went extinct. The phenotypic optimum was initially set to $\theta_0 = 0$, and held constant for 1,000 generations to allow the $V_{G,poly}$ to reach approximate selection-drift-mutation equilibrum before the onset of density-dependent population growth. We set up the simulations so that half of the repetitions would have a large effect, beneficial allele at relatively low frequency ($p_0 < 0.5$), and half would have a relatively high frequency ($p_0 > 0.5$) at the onset of density-dependent population growth and the environmentally-induced challenge to population persistence. To achieve this, we introduced an historical shift in the phenotypic optimum to allow an initially rare, large effect, beneficial allele to become relatively frequent before the onset of density dependent population growth (as seen in many natural populations, e.g., (Barson et al. 2015; Jones et al. 2018; Thompson et al. 2019)). Specifically, $\theta_0$ was shifted to $\theta_1$ in generation 1,001. We set $\theta_1 = 6$ in the 250 simulations with a desired $p_0 < 0.5$, and $\theta_1 = 12$ for the 250 simulations with desired $p_0 > 0.5$ (large historical shifts in $\theta$ result in greater change in allele freuquency). We initiated density-dependent population growth and increased $\theta$ by 10 units relative to the current population mean phenotype (for consistency with individual-based simulations described in prevsious sections) when the large effect allele was in the correct allele frequency range (i.e., $p < 0.5$ or $p > 0.5$), and $h^2$ was between 0.55 and 0.65 to control the $h^2$ at the onset of density-dependent population growth and for consistency with the previous individual-based simulations above. No mutations were allowed after the onset of density-dependent population growth in order to focus on evolutionary and population dynamics arising from standing genetic variation.

We then ran 1,000 additional simulations where the $V_G$ was due entirely to polygenic mutation ($\mu = 7.35 \times 10^{-8}$), such that $h^2 \approx 0.6$ at the onset of density-dependent population growth. Here, the simulations ran for 1,200 generations with $\theta_0 = 0$, and constant population size to allow the $V_G$ to reach approximate mutation-drift-selection equilibrium. We initiated density-dependent population growth and increased $\theta$ by 10 units relative to the current population mean phenotype in generation 1,201 (consisten with the simulations above with large-effect loci). We retained only 500 simulation repetitions where the realized $h^2$ was between 0.55 and 0.65 so that these simulations with a polygenic trait architecture were similar to the mutation-based simulations with large effect loci in terms of the $h^2$ at the onset of the environmentally-induced challenge to population persistence. We analyzed the output from the mutation-based simulations as for the individual-based simulations described in main text. the results from these simulations are shown in Figure S17.

### SUPPLEMENTARY FIGURES

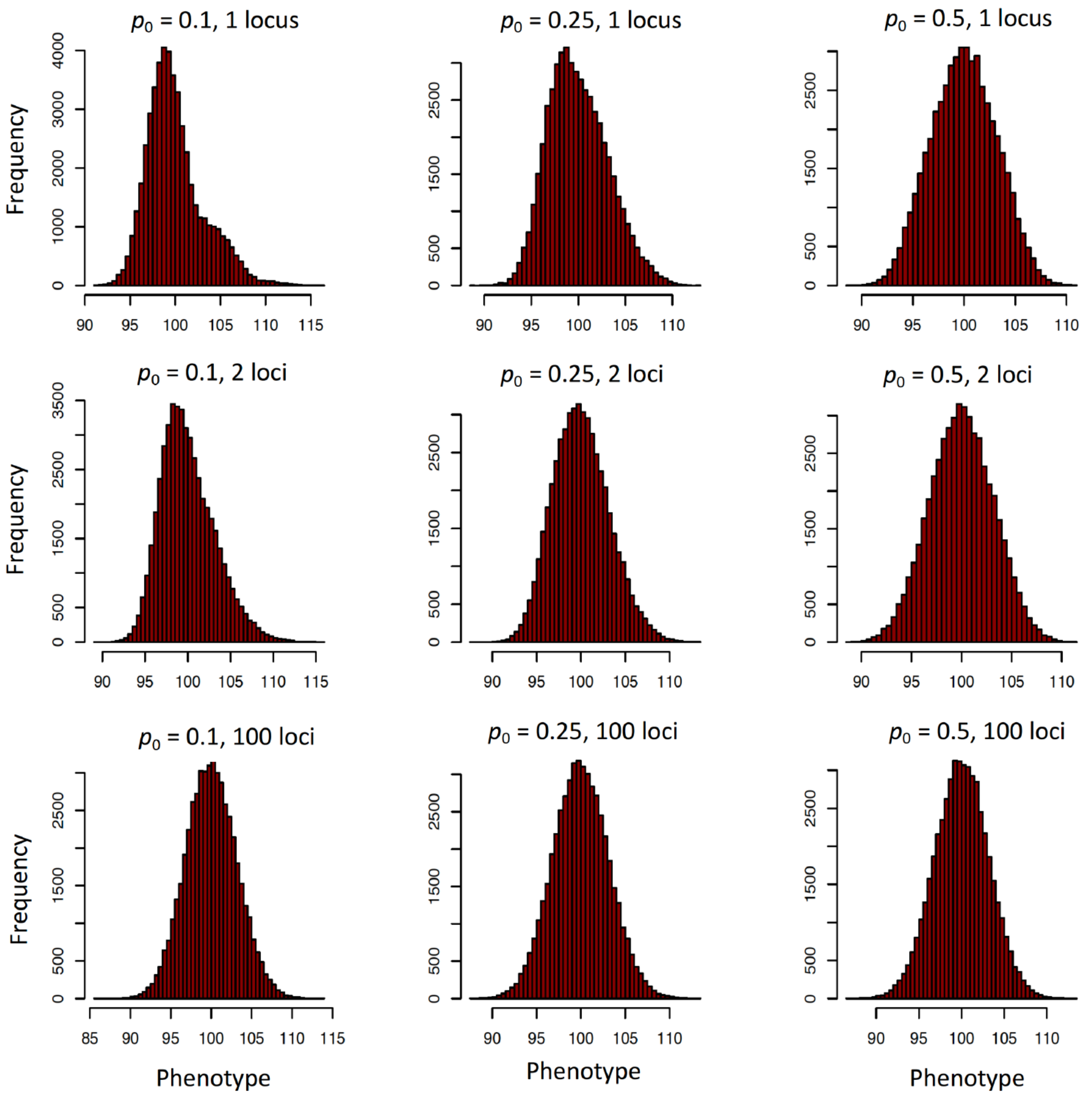


**Figure S1**. Distribution of simulated phenotypes. The simulated phenotype distributions are shown for three initial allele frequencies ($p_0$ = 0.1, 0.25, 0.5), and three numbers of loci contributing to the selected phenotype ($n$ = 1, 2, and 100 loci). Each panel shows the distribution of 50,000 simulated phenotypes.

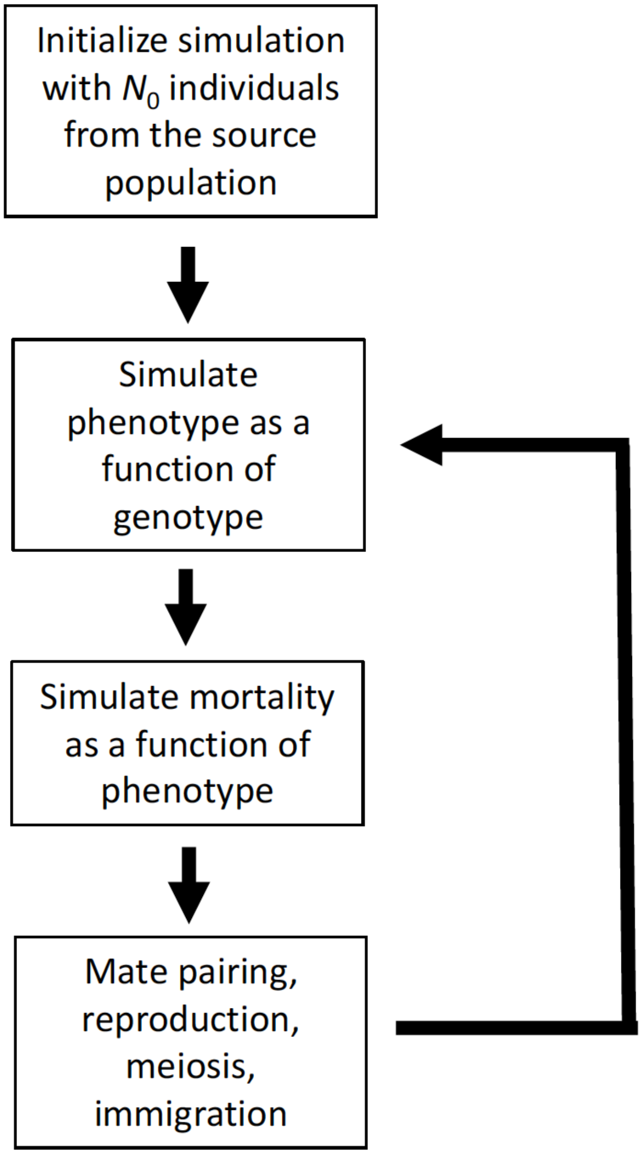


200 **Fig. S2.** Flowchart of the sequence of events in the simulation model. Details of each step in the simulation procedure are described in detail above in the Material and Methods.

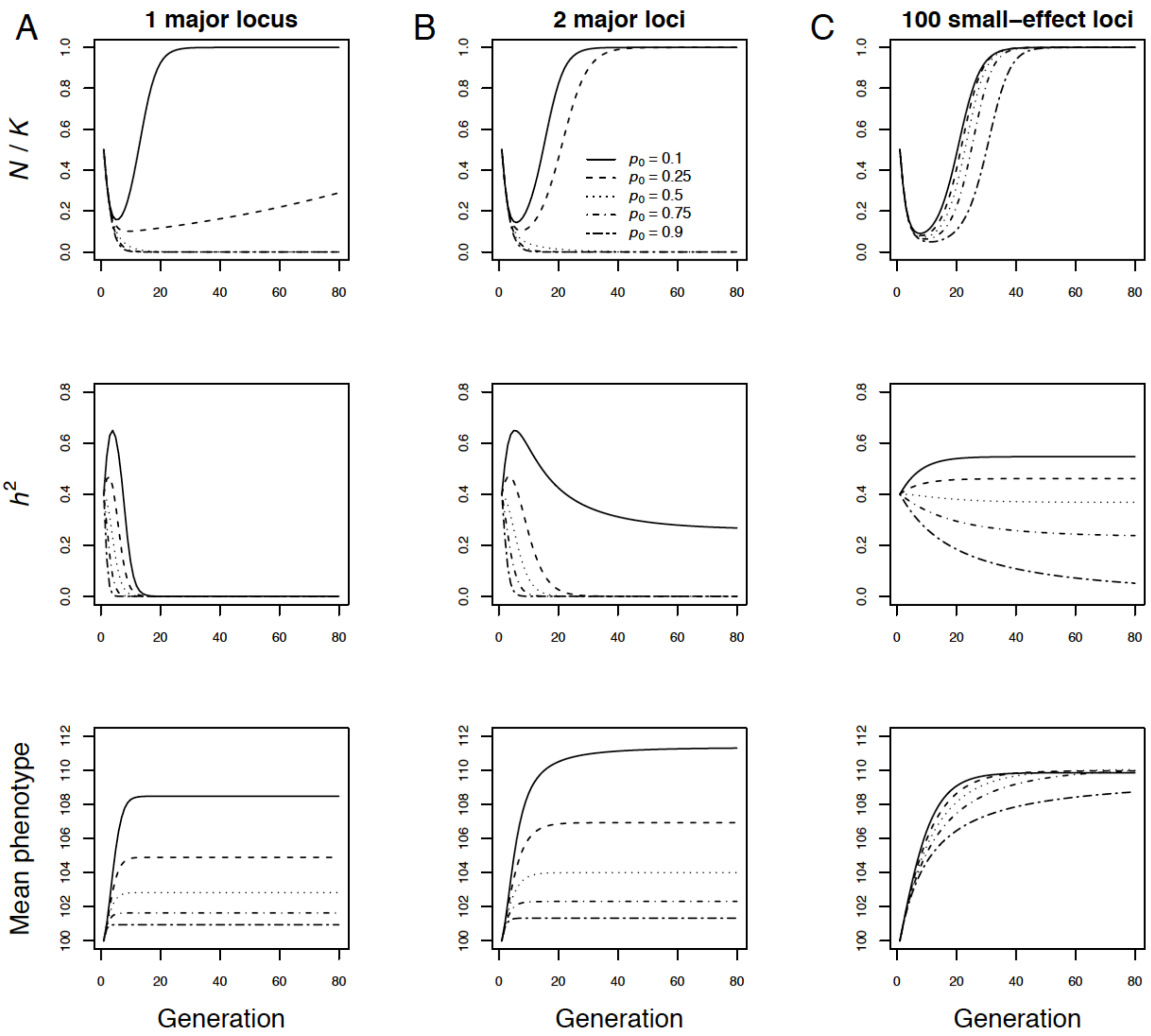


**Figure S3.** Deterministic predictions of evolutionary and demographic responses to directional selection in density-regulated populations with a single large-effect locus (**A**), two large-effect loci (**B**), and 100 small-effect loci (**C**) affecting a quantitative trait under directional selection after a sudden environmental change. The initial heritability was $h^2 = 0.4$ in all cases. Line types indicate the initial frequencies of the positively selected allele(s) conferring a larger phenotype. Initial population size was $K/2$.

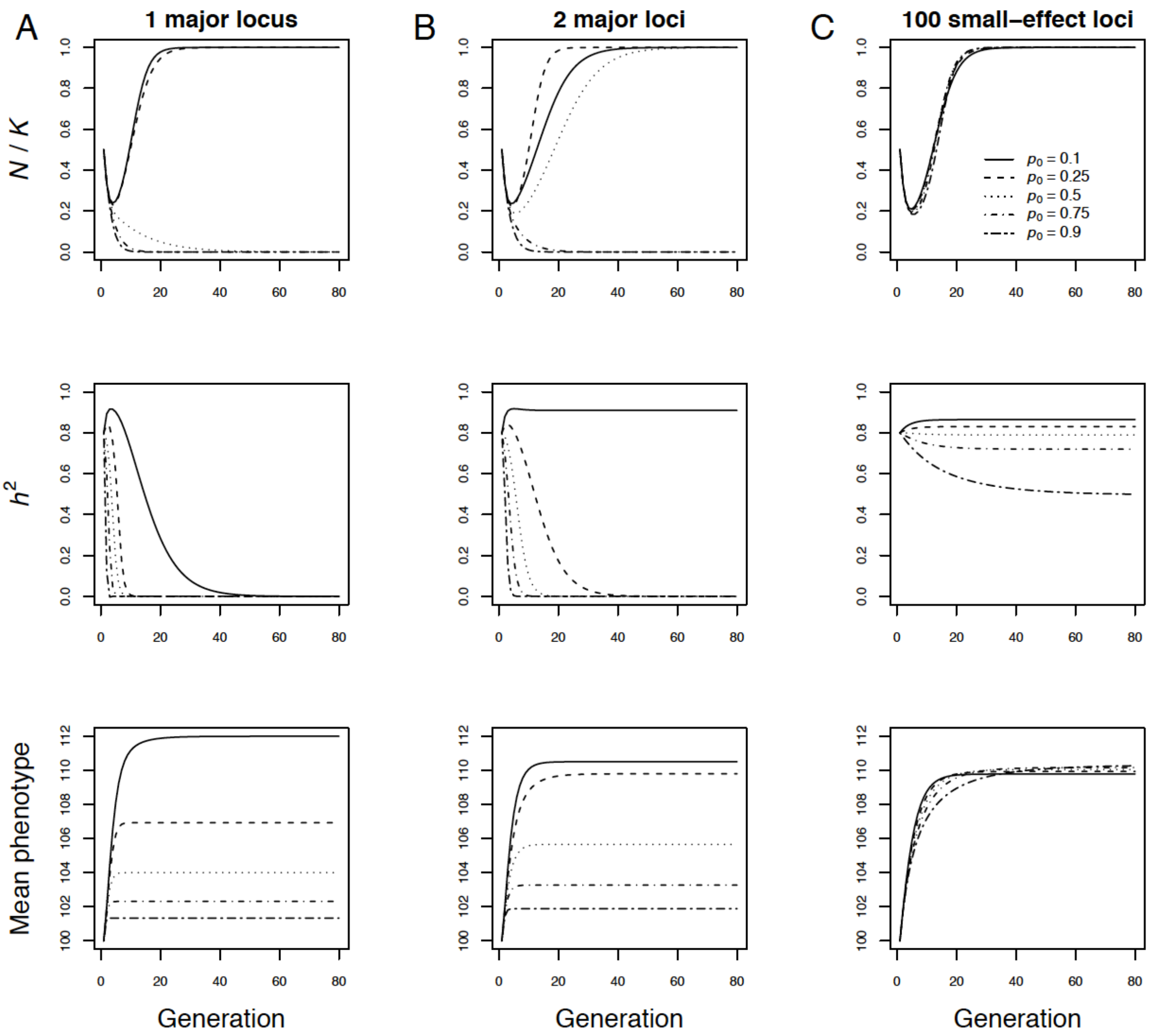


**Figure S4.** Deterministic predictions of evolutionary and demographic responses to directional selection in density-regulated populations with a single large-effect locus (**A**), two large-effect loci (**B**), and 100 small-effect loci (**C**) affecting a quantitative trait under directional selection after a sudden environmental change. The initial heritability was $h^2 = 0.8$ in all cases. Line types indicate the initial frequencies of the positively selected allele(s) conferring a larger phenotype. Initial population size was $K/2$.

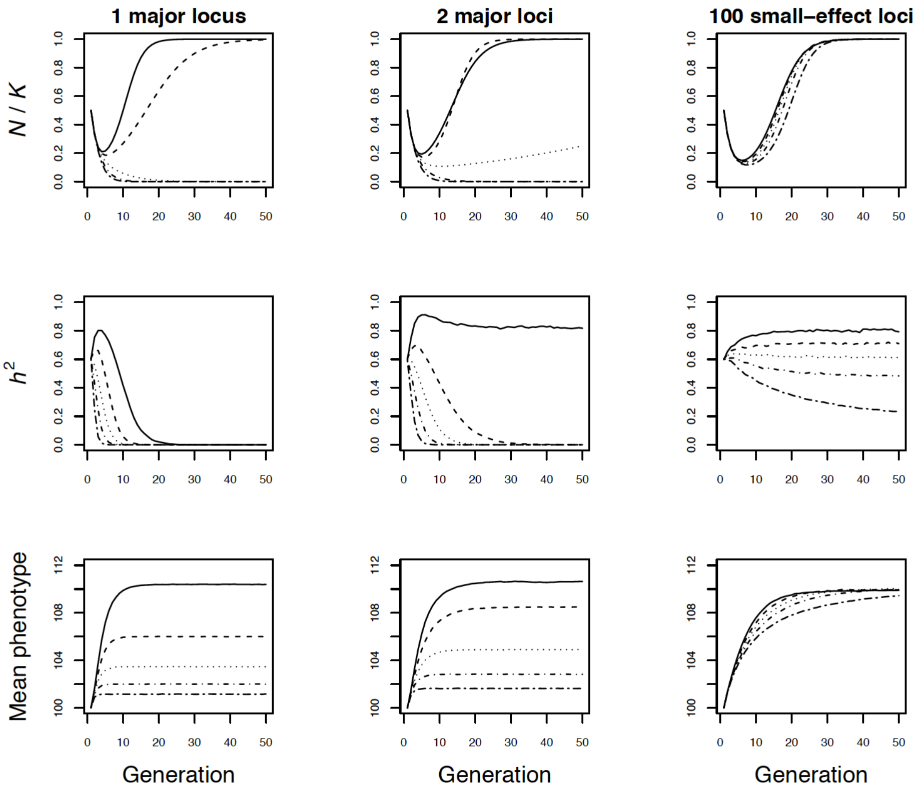


**Figure S5.** Evolutionary and demographic responses to a sudden change in the phenotypic optimum in density-regulated populations with a single large-effect locus (**A**), two large-effect loci (**B**), and 100 small-effect loci (**C**) affecting a quantitative trait under directional selection after a sudden environmental change. The lines in the first, second, and third rows show the mean population size, heritability, and mean phenotype across all 500 simulation repetitions versus time in generations. The initial heritability was $h^2 = 0.6$ in all cases. We varied the initial frequencies of the positively selected alleles conferring a larger phenotype from 0.1 to 0.9. Initial population size was $N_0 = 500$, and carrying capacity was $K$= 1,000 in all simulations. The lines show the mean of each parameter across 500 simulation repetitions. The mean $h^2$ and phenotype values shown were calculated across all populations with sizes > $N = 0$ each generation. The model used here is identical to that used to produce the data shown in Figure 1 in the main text, except here we simulated the distributions of individual genotypes and phenotypes to account for any selection-induced linkage disequilibrium and deviations from the normality in the phenotypic distribution. Specifically, each generation we simulated the genotypes, phenotypes, and mating among 50,000 pseudo individuals. The genotypes in the first generation were initialized using the assumed $p_0$. The phenotypes were simulated following the individual-based quantitative genetic model described in the main text. Offspring were assigned parents using weighted sampling (with the *sample* function in R), with the weights assigned based on phenotype using equation (1) in the main text. Population size in year t + 1 ($N_{t+1}$) was determined with equation (10) in the main text, with $\overline{w}_t$ replaced with the mean fitness among the 50,000 pseudo individuals in year $t$. This approach allowed us to precisely determine the expected distribution of individual fitness while accounting for any selection-induced linkage disequilibrium and non-normality in the phenotype distribution.

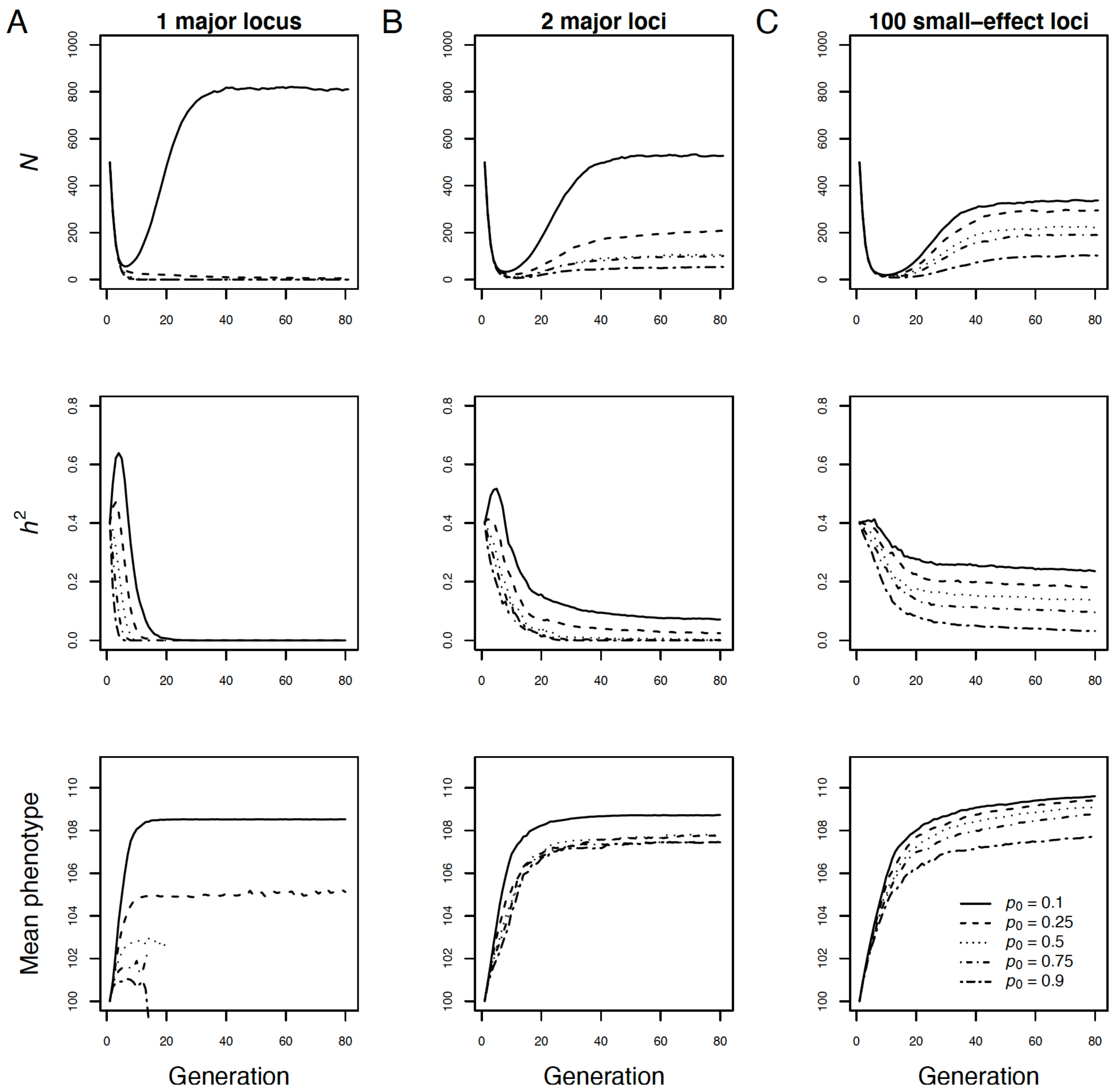


**Figure S6.** Individual-based simulations of evolutionary and demographic responses to directional selection in density-regulated populations with a single large-effect locus (**A**), two large-effect loci (**B**), and 100 small-effect loci (**C**) affecting a quantitative trait under directional selection after a sudden environmental change. The initial heritability was $h_0^2 = 0.4$ in all cases. Line types indicate the initial frequencies of the positively selected allele(s) conferring a larger phenotype. Initial population size was $N_0 = 500$, and varying capacity was $K = 1{,}000$. The extinction rates from these simulations are shown in Figure S6. The lines in the first, second, and third rows show the mean population size, heritability, and mean phenotype across all 500 simulation repetitions versus time in generations.

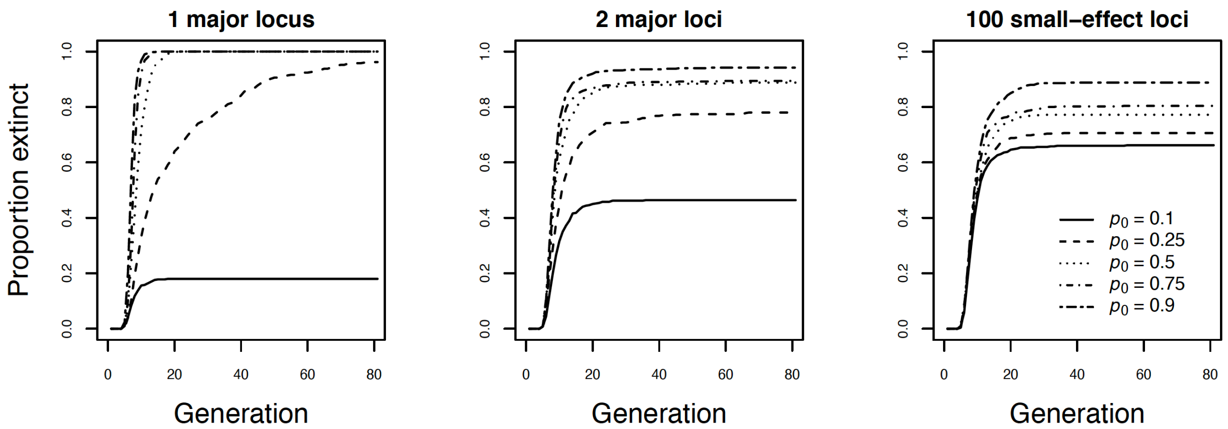


**Figure S7.** Extinction rates in density-regulated populations with a single large-effect locus (**A**), two large-effect loci (**B**), and 100 small-effect loci (**C**) affecting a quantitative trait under directional selection after a sudden environmental change. The initial heritability was $h_0^2 = 0.4$ in all cases. Line types indicate the initial frequencies of the positively selected allele(s) conferring a larger phenotype. Initial population size was $N_0 = 500$, and varying capacity was $K = 1{,}000$.

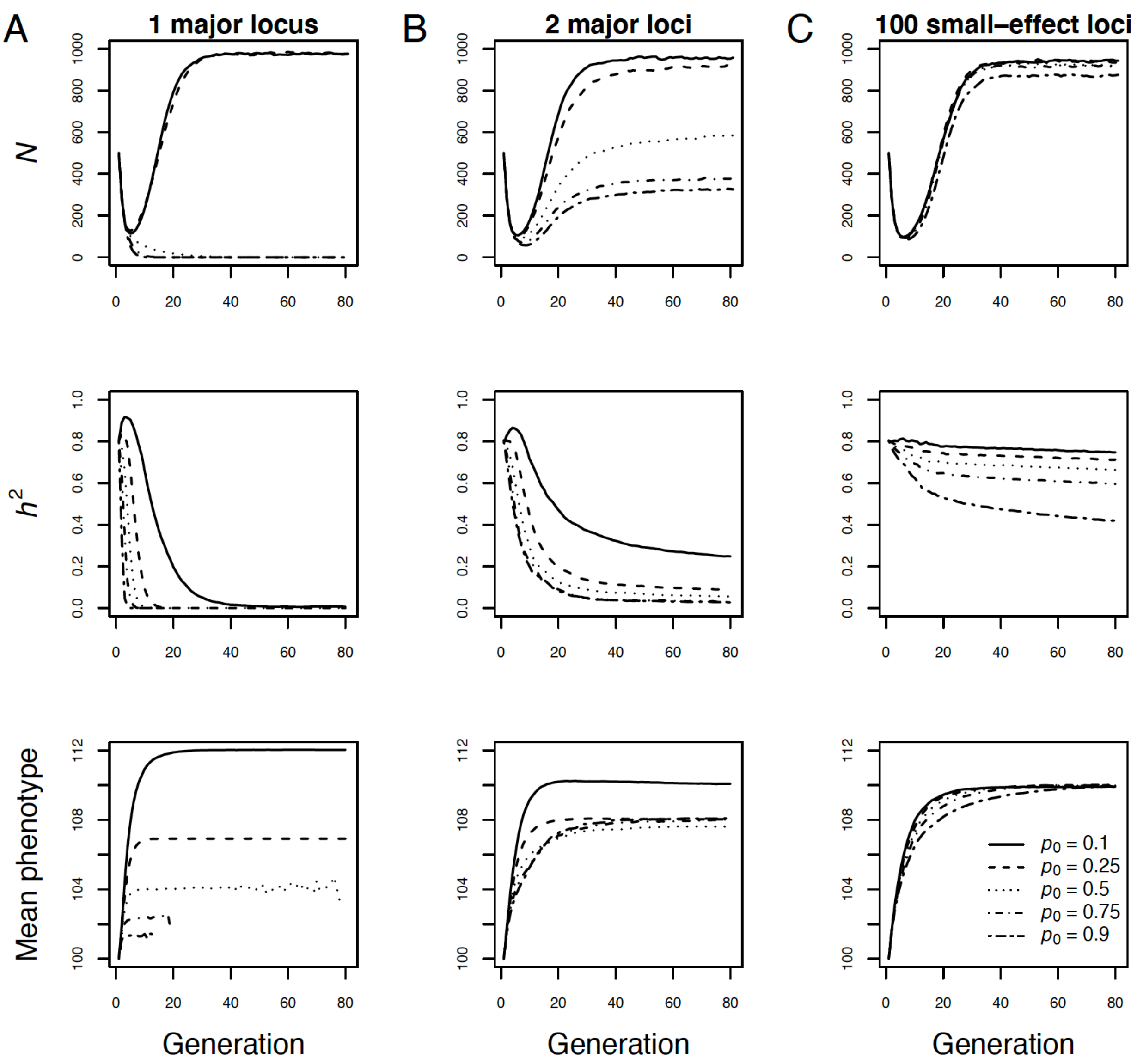


**Figure S8.** Individual-based simulations of evolutionary and demographic responses to directional selection in density-regulated populations with a single large-effect locus (**A**), two large-effect loci (**B**), and 100 small-effect loci (**C**) affecting a quantitative trait under directional selection after a sudden environmental change. The lines in the first, second, and third rows show the mean population size, heritability, and mean phenotype across all 500 simulation repetitions versus time in generations. The initial heritability was $h_0^2 = 0.8$ in all cases. Line types indicate the initial frequencies of the positively selected allele(s) conferring a larger phenotype. Initial population size was $N_0 = 500$, and varying capacity was $K = 1{,}000$. The extinction rates from these simulations are shown in Figure S8.

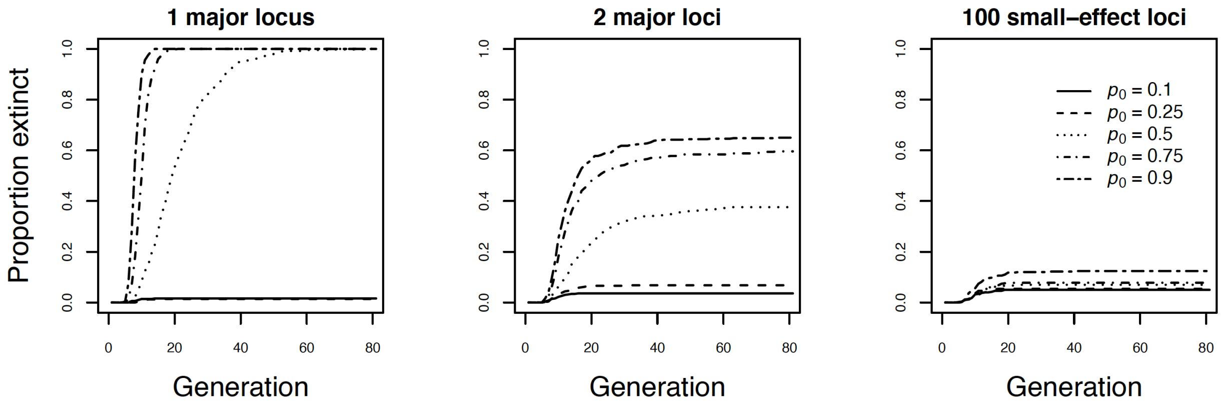


**Figure S9.** Extinction rates in density-regulated populations with a single large-effect locus (**A**), two large-effect loci (**B**), and 100 small-effect loci (**C**) affecting a quantitative trait under directional selection after a sudden environmental change. The initial heritability was $h_0^2 = 0.8$ in all cases. Line types indicate the initial frequencies of the positively selected allele(s) conferring a larger phenotype. Initial population size was $N_0 = 500$, and varying capacity was $K = 1{,}000$.

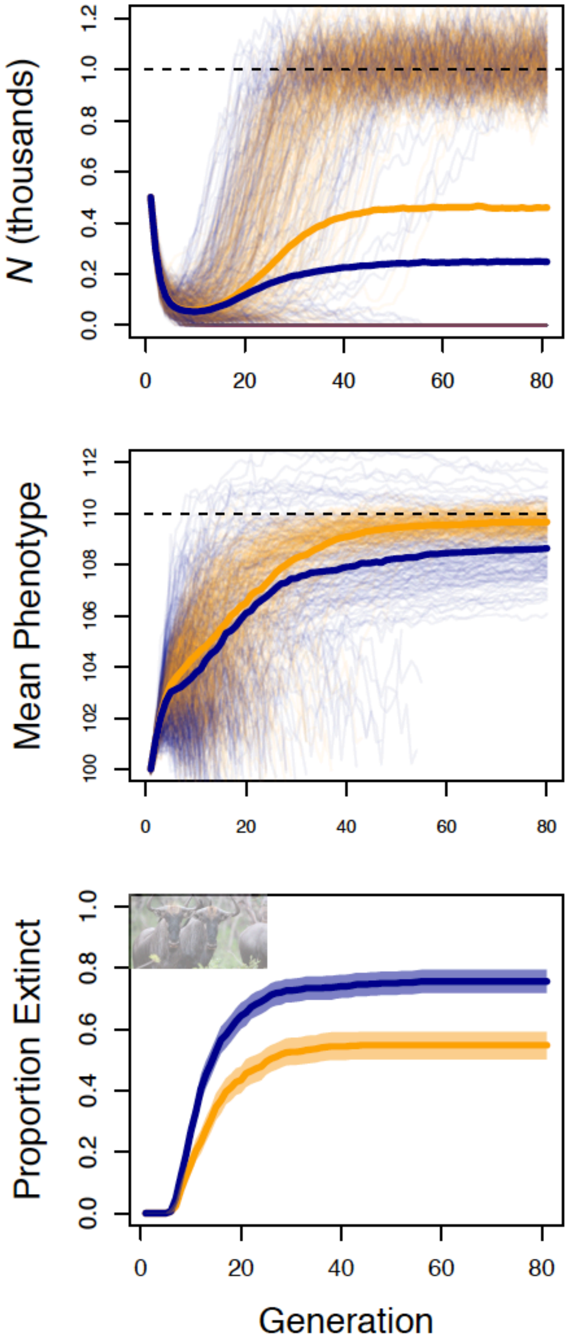


**Figure S10.** Effects of genetic architecture on phenotypic evolution and population dynamics in populations with an approximate large mammal life history and a low immigration rate (4 immigrants per generation) from a population with a constant phenotypic optimum of size = 100. Results from populations with a large-effect locus are shown in blue; populations where the selected trait was polygenic are in orange. Thin lines show the population size (top row) and mean phenotype (middle row) through time. Thick lines show the mean population size and phenotype across all 500 repetitions. The bottom panels show the proportion of extinct populations through time, with bootstrap 95% confidence intervals.

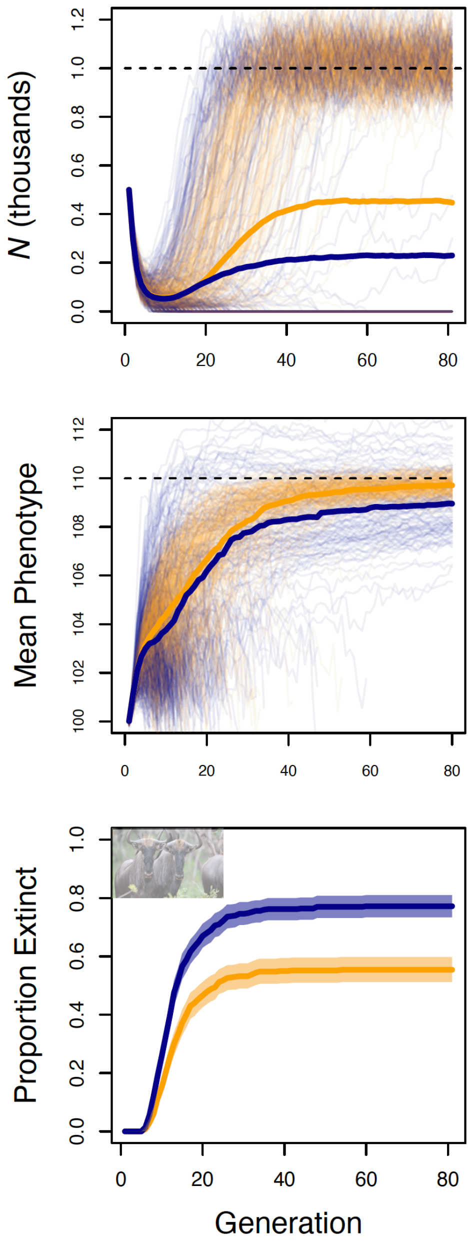


**Figure S11.** Effects of genetic architecture on phenotypic evolution and population dynamics in populations with an approximate large mammal life history and a high immigration rate (8 immigrants per generation) from a population with a constant phenotypic optimum of size = 100. Results from populations with a large-effect locus are shown in blue; populations where the selected trait was polygenic are in orange. Thin lines show the population size (top row) and mean phenotype (middle row) through time. Thick lines show the mean population size and phenotype across all 500 repetitions. The bottom panels show the proportion of extinct populations through time, with bootstrap 95% confidence intervals.

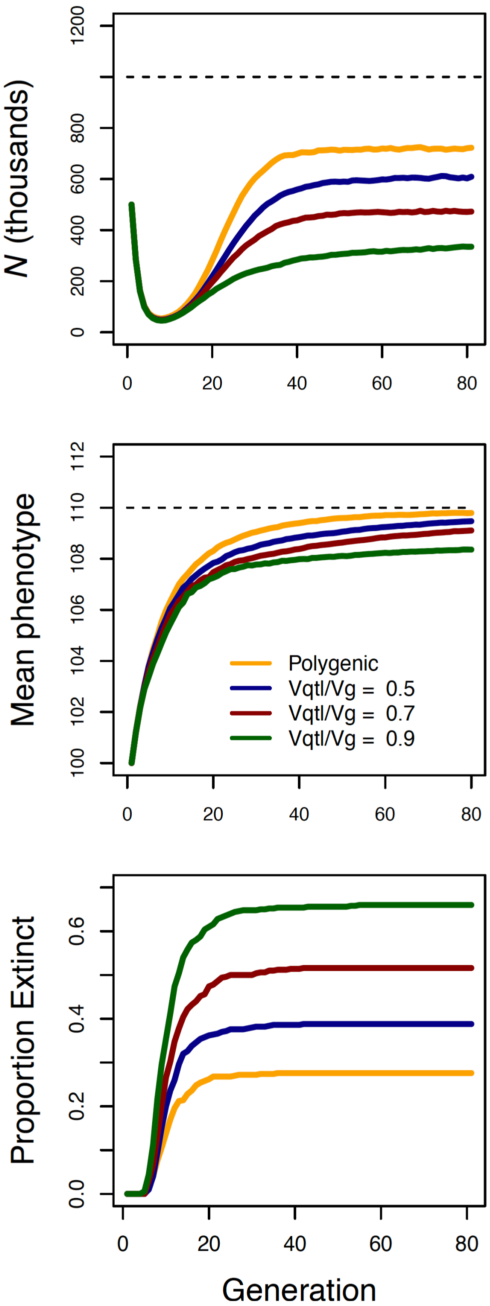


**Figure S12.** Effects of the effect size of large-effect loci on population dynamics.
The results are from simulations parameterized as in our simulations shown in Figure 3A, except here we varied the proportion of the genetic variance attributed to the large effect locus. Results from populations where a major locus was responsible for 90% (green), 70% (red), 50% (blue) of the additive genetic variance ($V_G$). Orange lines show results from populations where there selected trait was polygenic (no large-effect locus). The bottom panel shows the proportion of extinct populations though time.

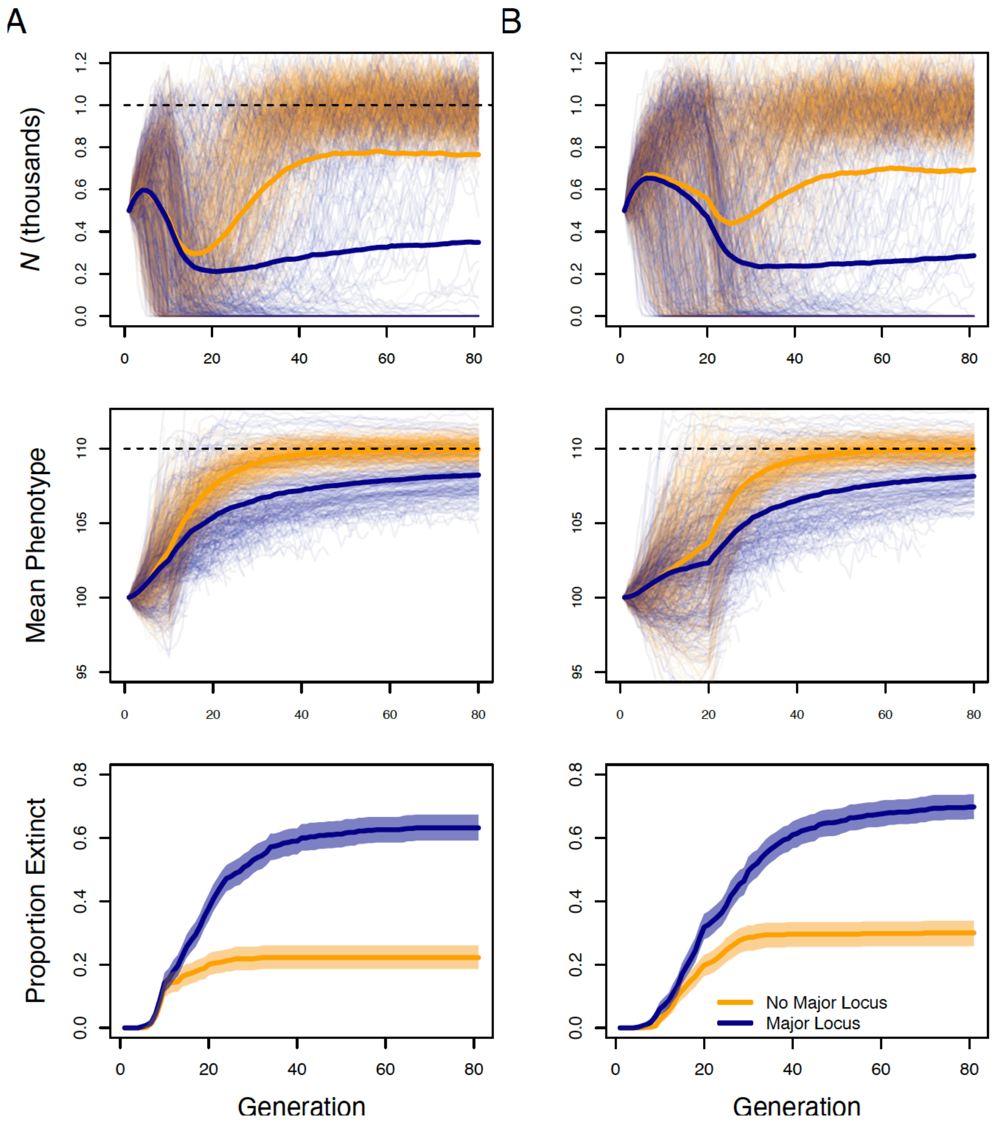


**Figure S13.** Effects of genetic architecture on phenotypic evolution and population dynamics in populations with a large mammal-like life history and with a stochastic linear increase in the optimum phenotype value with time. We ran simulations where the expected optimum phenotype increased linearly from 100 to 110 in either 10 (**A**) or 20 (**B**) generatiosn. Results from populations with a large-effect locus are shown in blue; populations where the selected trait was polygenic are in orange. Thin lines show the population size (top row) and mean phenotype (middle row) through time. Thick lines show the mean population size and phenotype across all 500 repetitions. The bottom panels show the proportion of extinct populations through time, with bootstrap 95% confidence intervals.

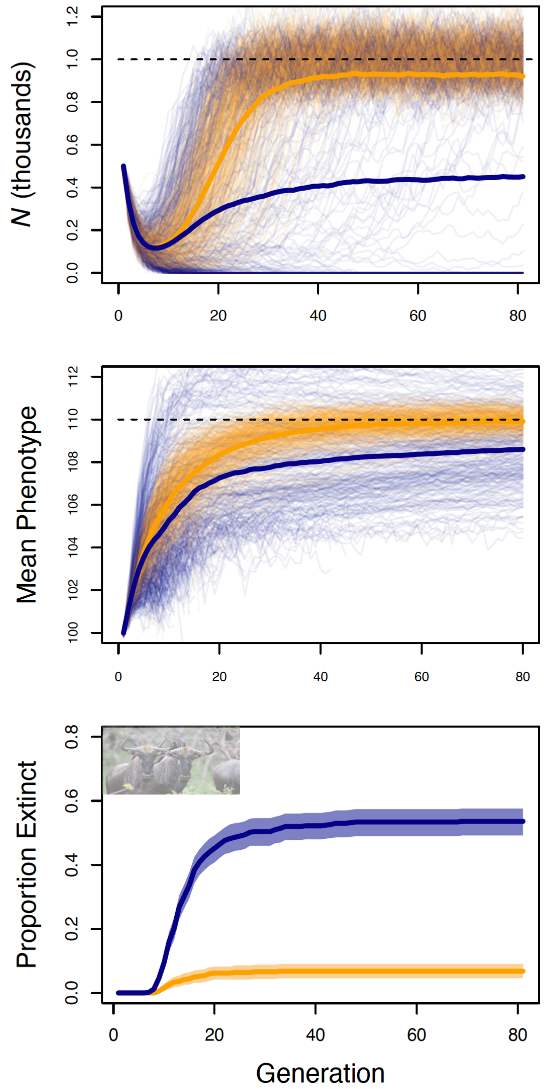


**Figure S14.** Effects of genetic architecture on phenotypic evolution and population dynamics with linkage in closed populations with a life history approximating large mammals. Results from populations with a large-effect locus are shown in blue; populations where the selected trait was polygenic are in orange. Thin lines show the population size (top row) and mean phenotype (middle row) through time. Thick lines show the mean population size across all simulation replicates. The bottom panels show the proportion of extinct populations through time, with bootstrap 95% confidence intervals.

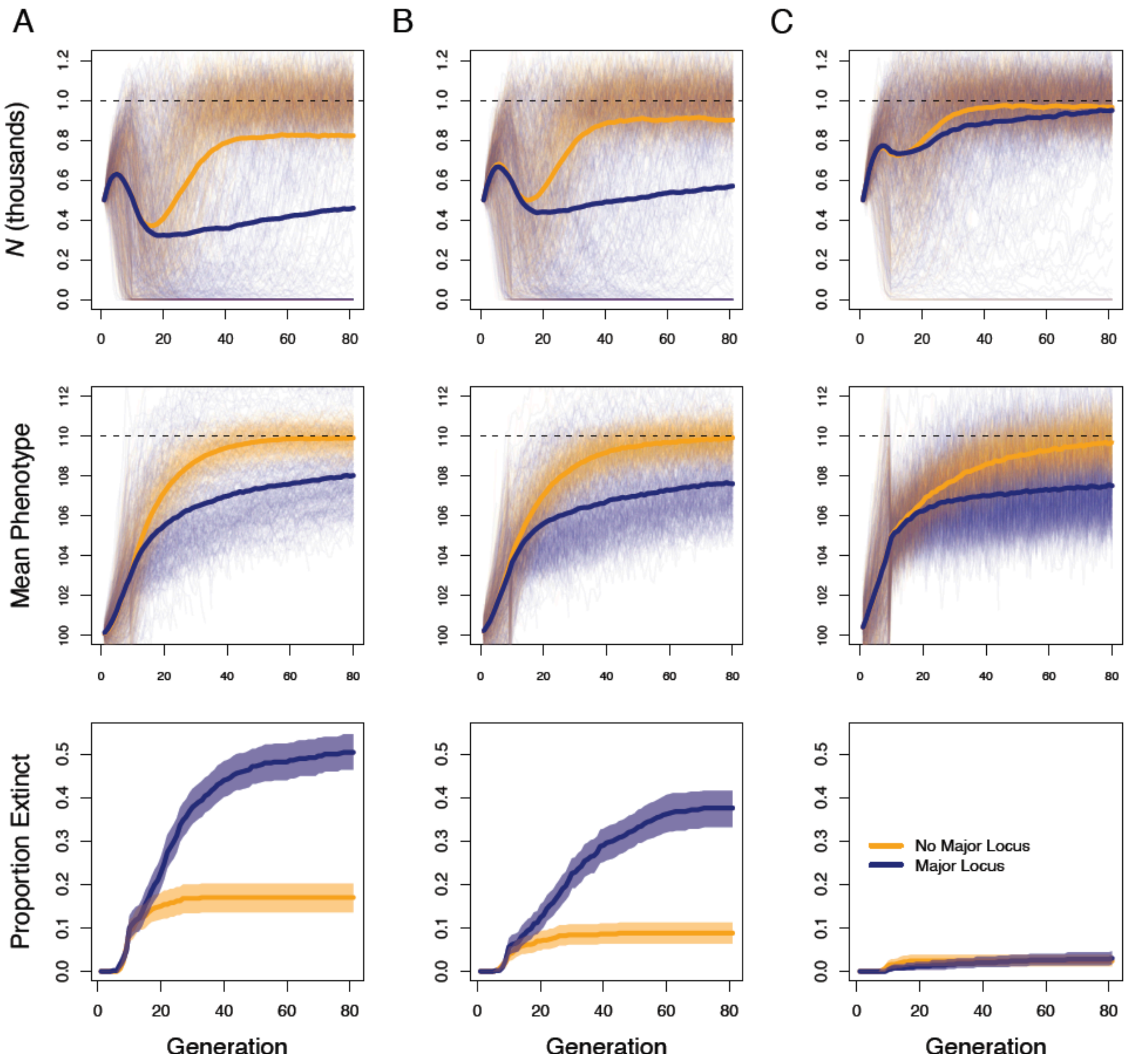


**Figure S15.** Effects of genetic architecture on phenotypic evolution and population dynamics in populations with life history approximating large mammals and plasticity in the selected phenotype. Results are shown form simulations with the strength of selection set to $m = 0.1$ (**A**), $m = 0.2$ (**B**), $m = 0.4$ (**C**). Results from populations with a large-effect locus are shown in blue; populations where the selected trait was polygenic are in orange. Thin lines show the population size (top row) and mean phenotype (middle row) through time. Thick lines show the mean population size across all simulation replicates. The bottom panels show the proportion of extinct populations through time, with bootstrap 95% confidence intervals.

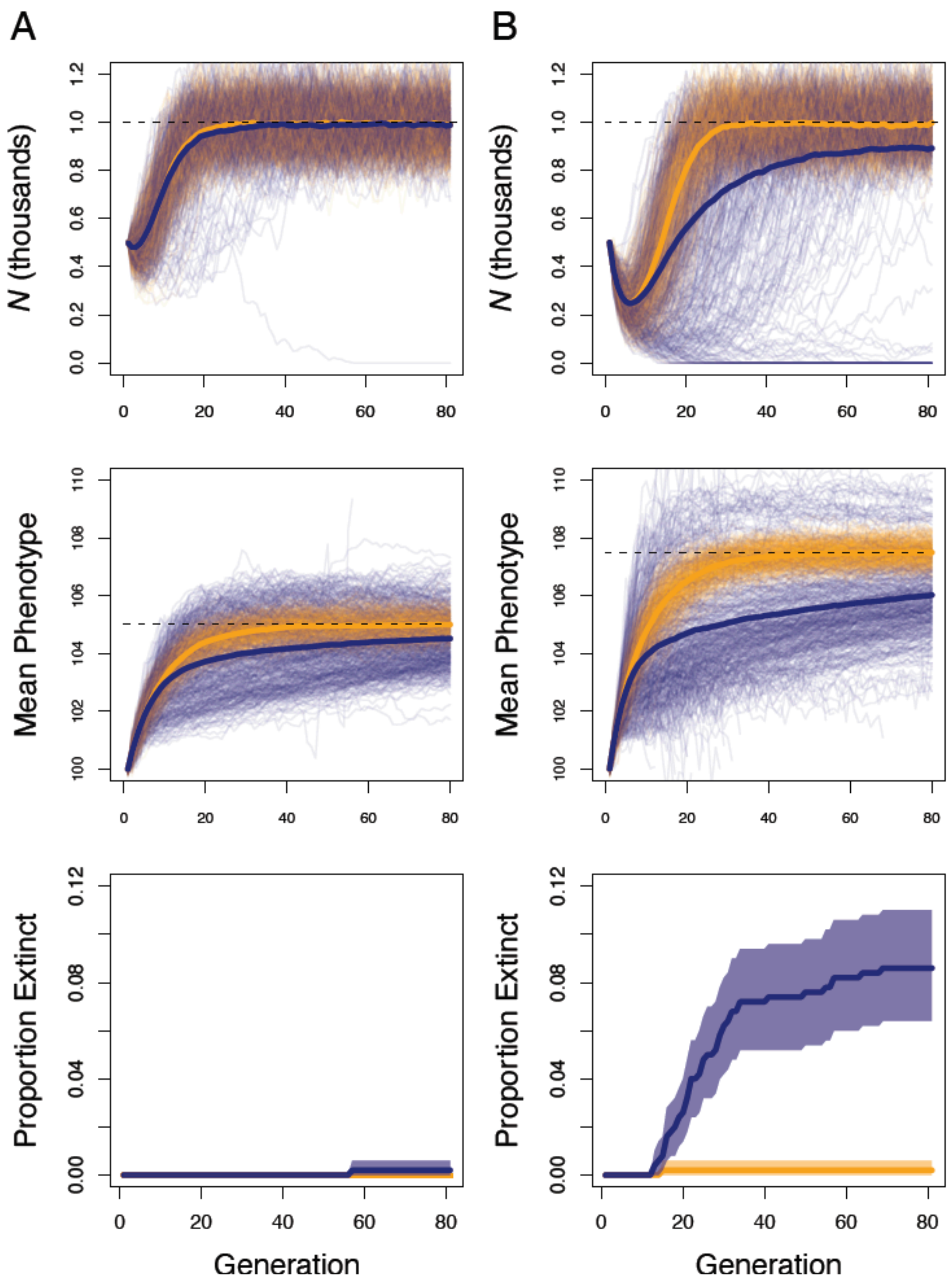


**Figure S16.** Effects of genetic architecture on phenotypic evolution and population dynamics in populations with life history approximating large mammals and plasticity in the selected phenotype. Results are shown form simulations equivalent to those shown in Figure 4 in the main text, except here the phenotypic optimum $\theta$ shifted from 100 to either 105 (**A**) or 107.5 (**B**) instead of to 110. Results from populations with a large-effect locus are shown in blue; populations where the selected trait was polygenic are in orange. Thin lines show the population size (top row) and mean phenotype (middle row) through time. Thick lines show the mean population size across all simulation replicates. The bottom panels show the proportion of extinct populations through time, with bootstrap 95% confidence intervals.

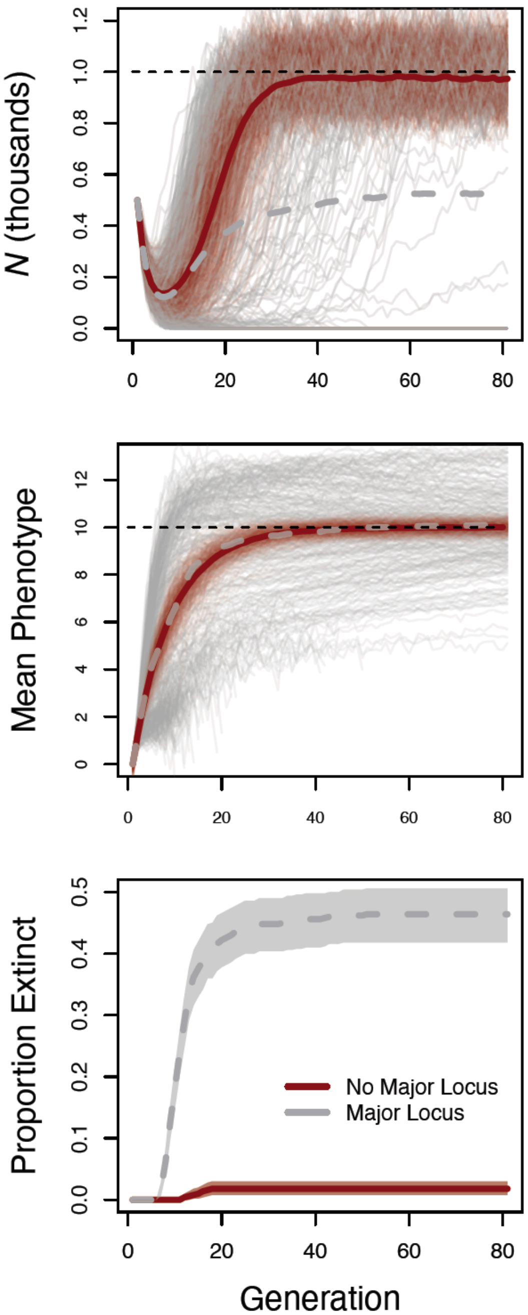


**Figure S17.** Effects of genetic architecture on phenotypic evolution and population dynamics in populations with life histories approximating large mammals and plasticity in the selected phenotype. Results are shown from simulations with a long burn and mutation to allow the genetic variance to reach approximate equilibrium. Results from populations with a large-effect locus are shown in gray; populations where the selected trait was polygenic are in dark red. Thin lines show the population size (top panel) and mean phenotype (middle panel) through time for each of 500 simulation repetitions. Thick lines show the mean population size and phenotype across all simulation replicates. The bottom panel shows the proportion of extinct populations through time, with bootstrap 95% bootstrap confidence intervals.

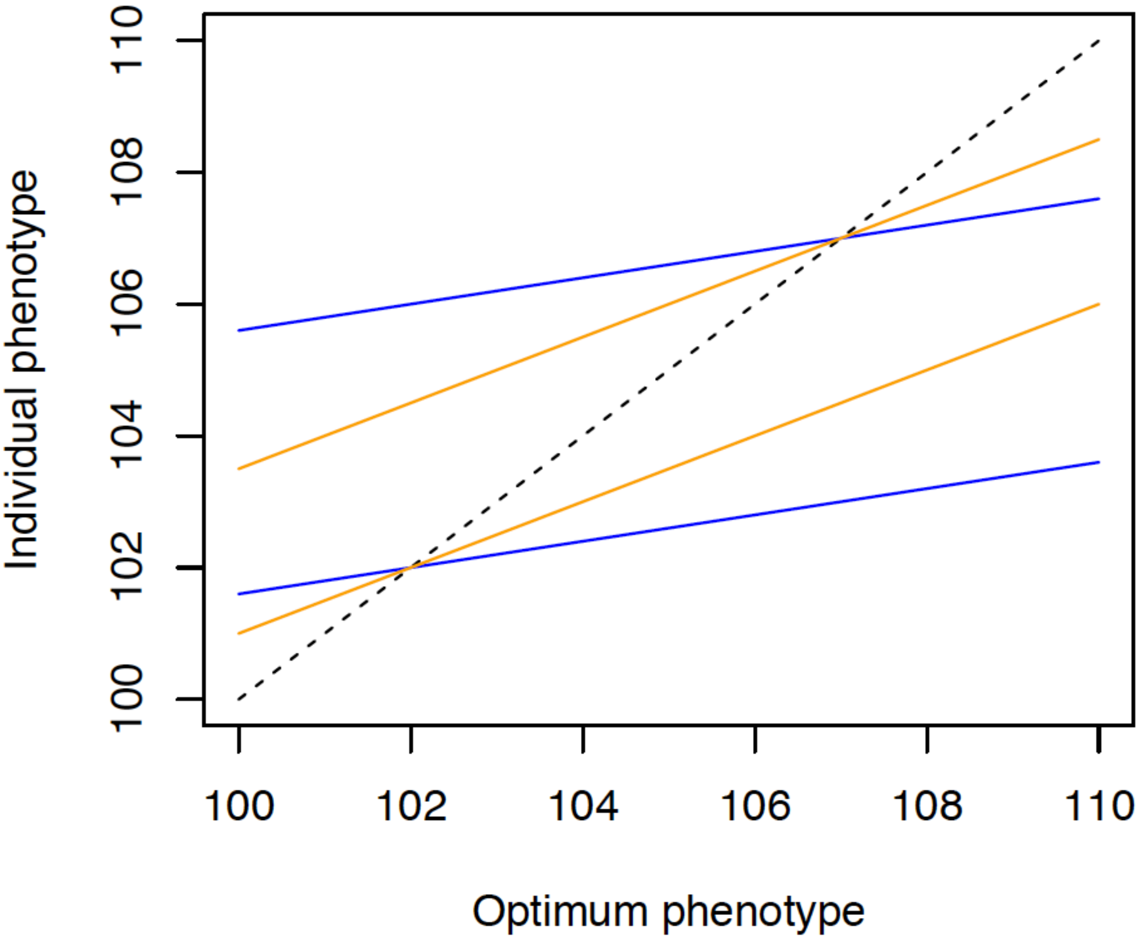


**Figure S18.** Model of plasticity shown as the individual phenotype plotted against the optimum phenotype. The dashed black line represents the case where individual phenotype equals the optimum phenotype. The expected phenotypes (i.e., ignoring random environmental effects) are shown for two individuals with breeding values of 102 and 107. The blue lines show the expected phenotypes with plasticity parameter *m* = 0.2, and the orange lines show the expected phenotypes with *m* = 0.5.

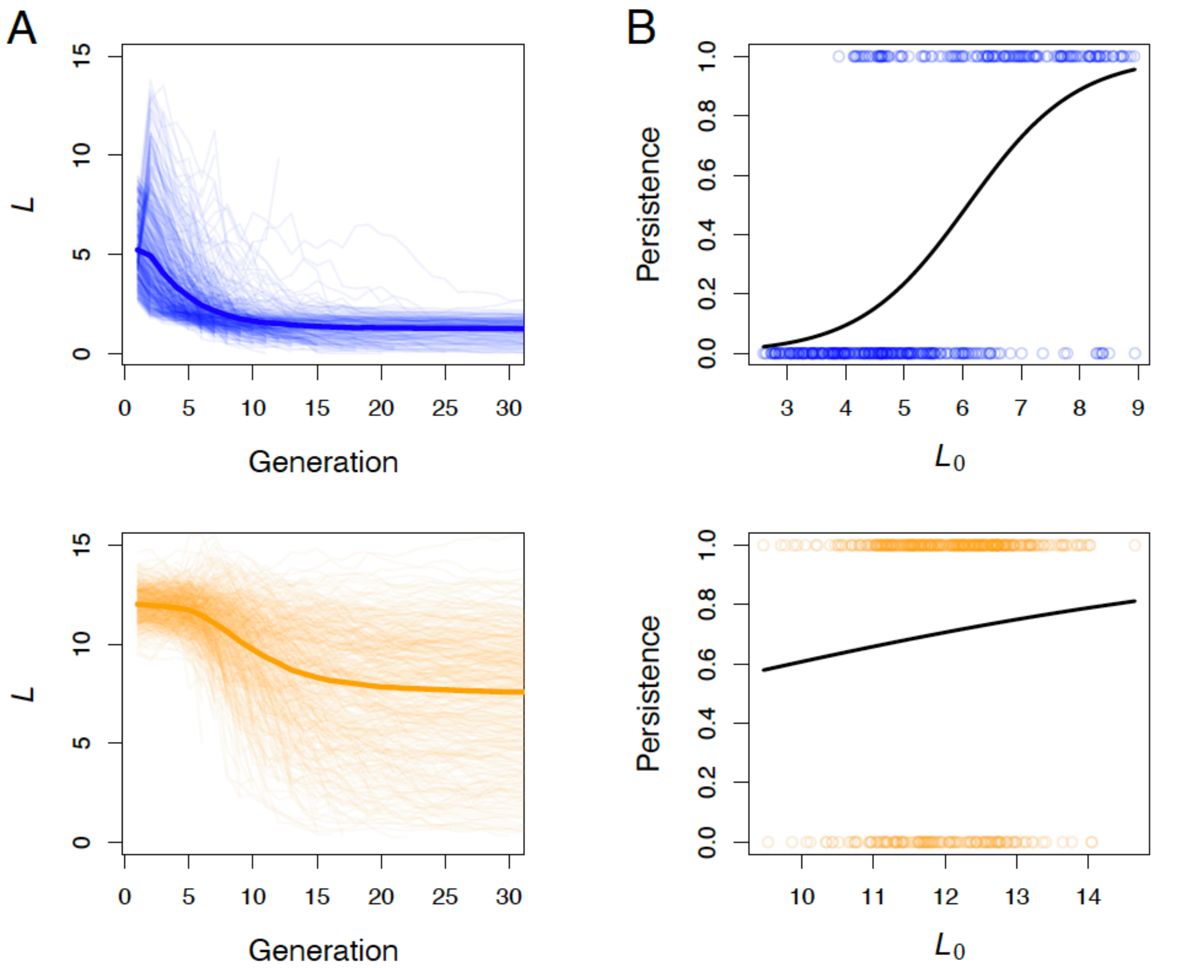


**Figure S19.** Temporal dynamics of the selection limit ($L$)(**A**), and the relationship between population viability and the selection limit at the beginning of the simulations ($L_0$) (**B**). Results are shown for simulated populations with a major locus responsible for 90% of $V_G$ (top row, blue lines), and where the selected trait was polygenic (bottom row, orange lines). The black lines in **B** are fitted logistic regression lines.

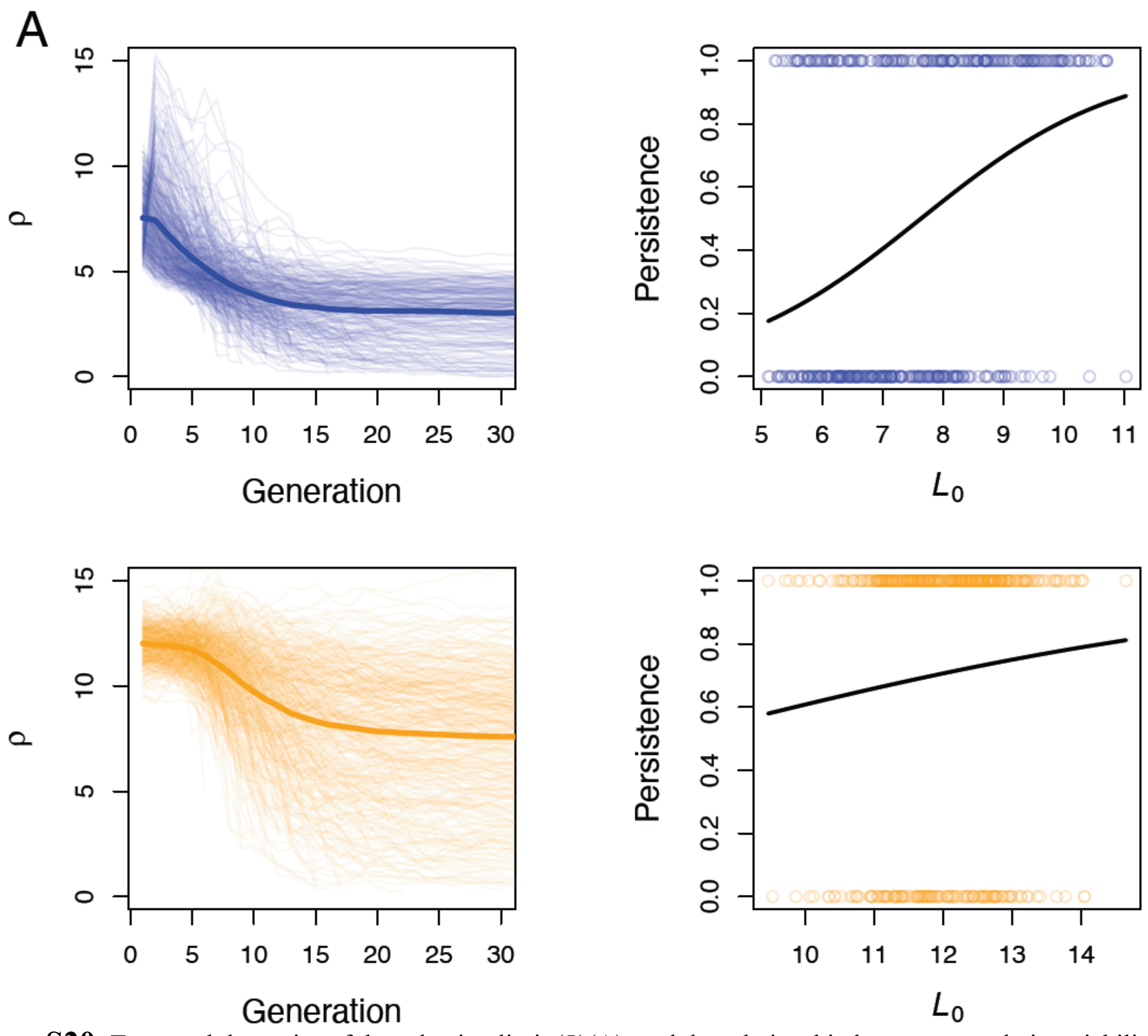


**Figure S20.** Temporal dynamics of the selection limit ($L$)(**A**), and the relationship between population viability and the selection limit at the beginning of the simulations ($L_0$) (**B**). Results are shown for simulated populations with a major locus responsible for 70% of $V_G$ (top row, blue lines), and where the selected trait was polygenic (bottom row, orange lines). The black lines in **B** are fitted logistic regression lines.

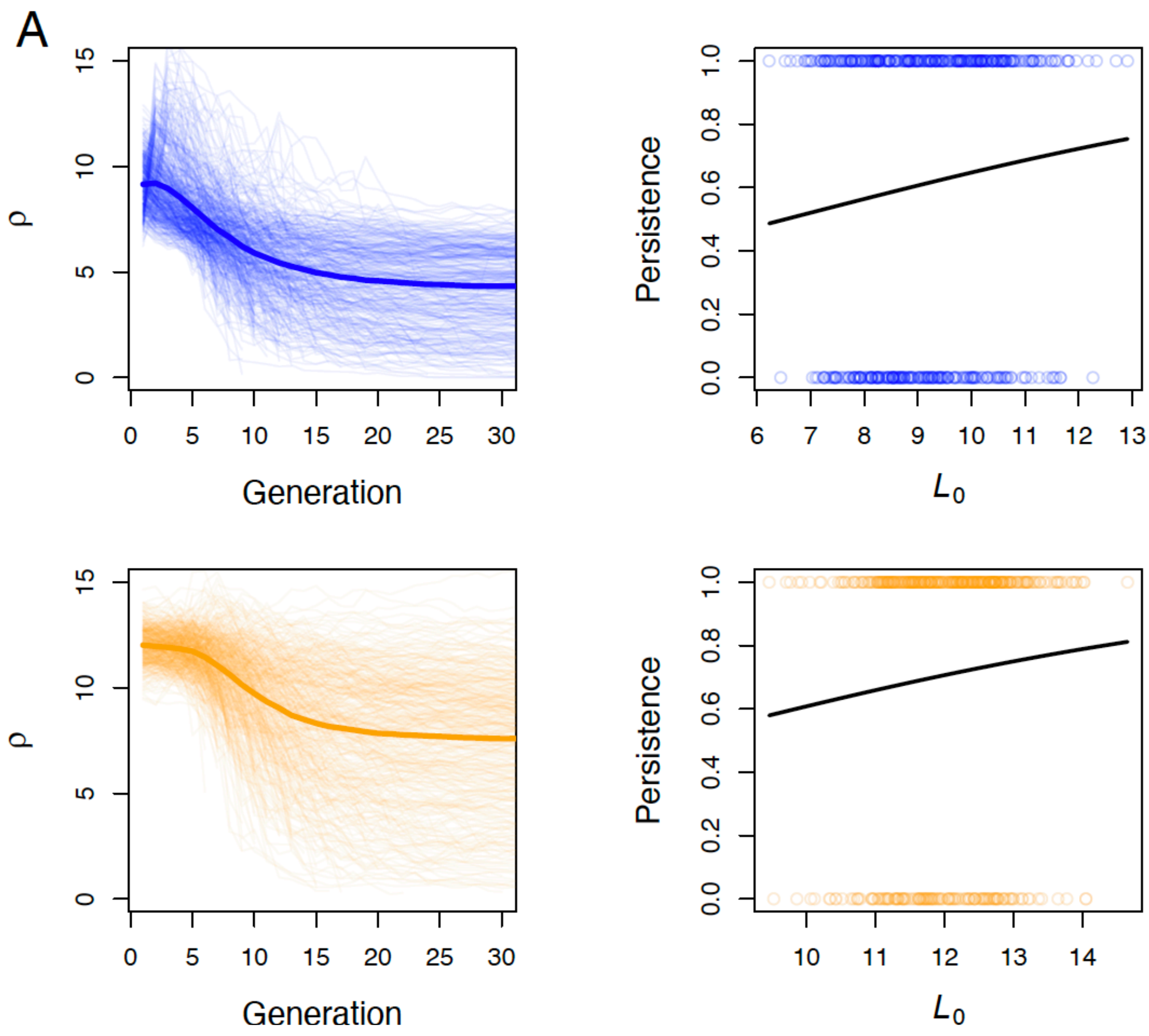


**Figure S21.** Temporal dynamics of the selection limit (*L*)(**A**), and the relationship between population viability and the selection limit at the beginning of the simulations ($L_0$) (**B**). Results are shown for simulated populations with a major locus responsible for 50% of $V_G$ (top row, blue lines), and where the selected trait was polygenic (bottom row, orange lines). The black lines in **B** are fitted logistic regression lines.

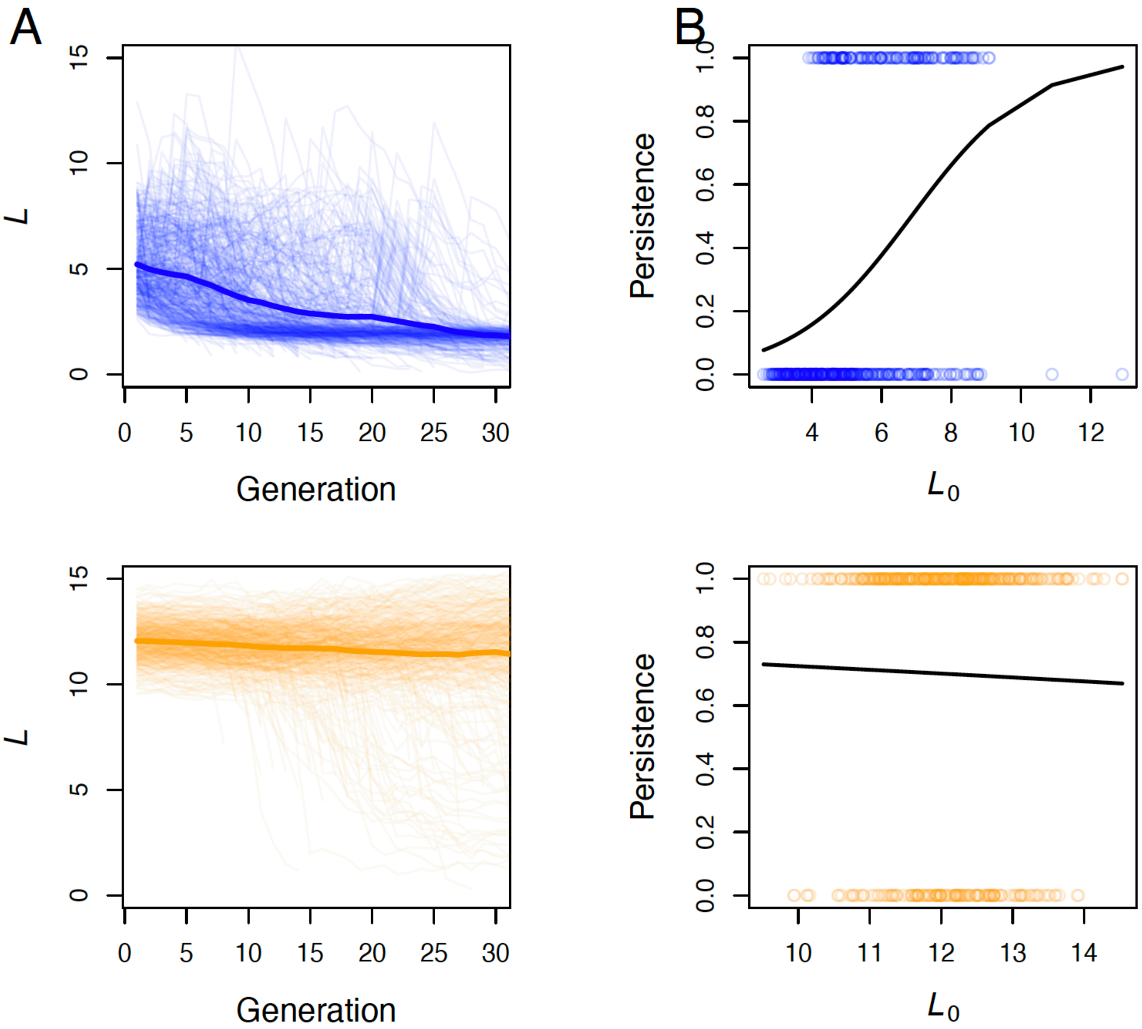


**Figure S22.** Temporal dynamics of the selection limit ($L$)(**A**), and the relationship between population viability and the selection limit at the beginning of the simulations ($L_0$) (**B**). Results are shown for simulated populations with a major locus responsible for 90% of $V_G$ (top row, blue lines), and where the selected trait was polygenic (bottom row, orange lines). The optimum phenotype in these simulations shifted from 100 to 110 over the first 20 generations. The black lines in **B** are fitted logistic regression lines.

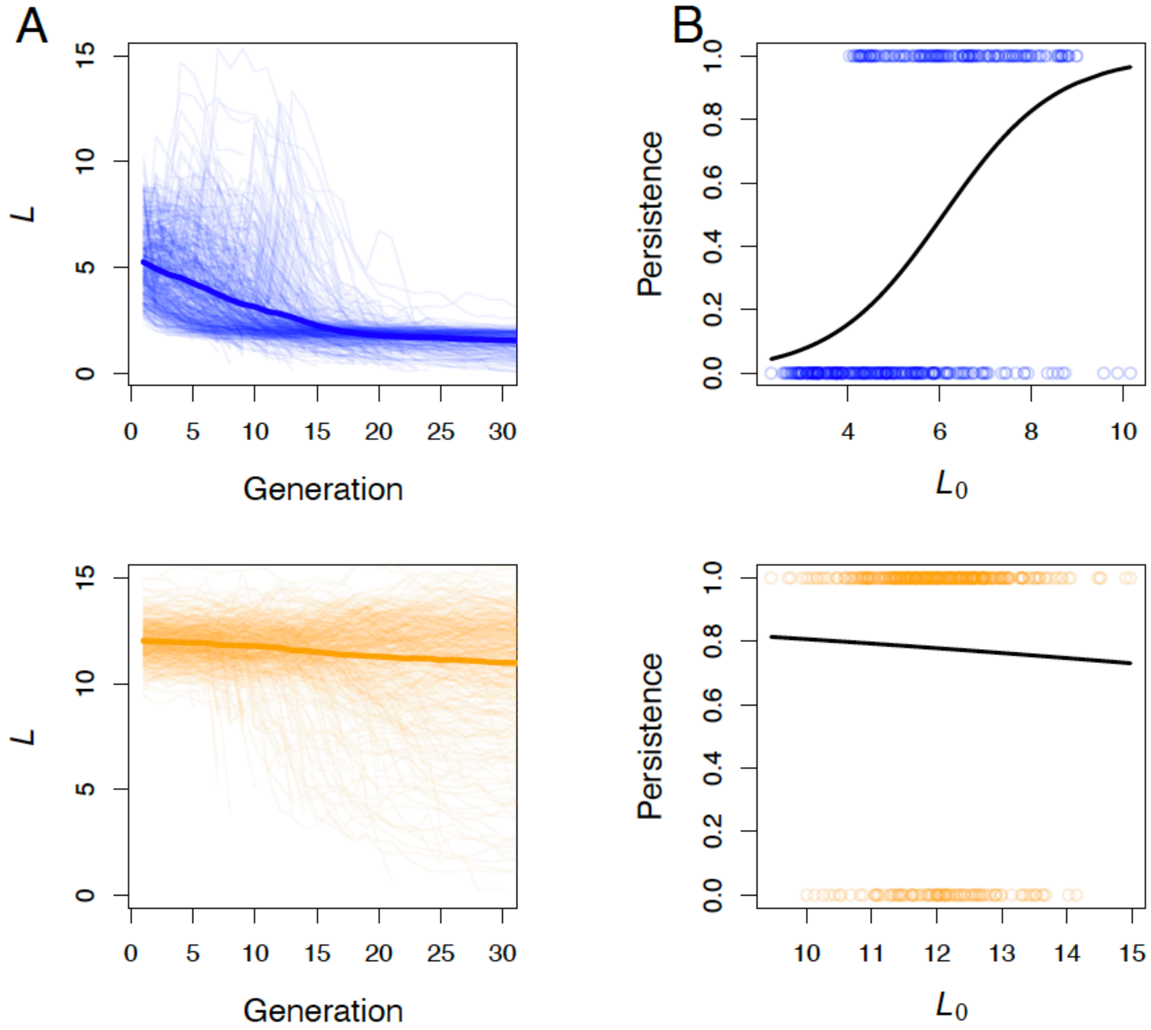


**Figure S23.** Temporal dynamics of the selection limit ($L$)(**A**), and the relationship between population viability and the selection limit at the beginning of the simulations ($L_0$) (**B**). Results are shown for simulated populations with a major locus responsible for 90% of $V_G$ (top row, blue lines), and where the selected trait was polygenic (bottom row, orange lines). The optimum phenotype in these simulations shifted from 100 to 110 over the first 10 generations. The black lines in **B** are fitted logistic regression lines.

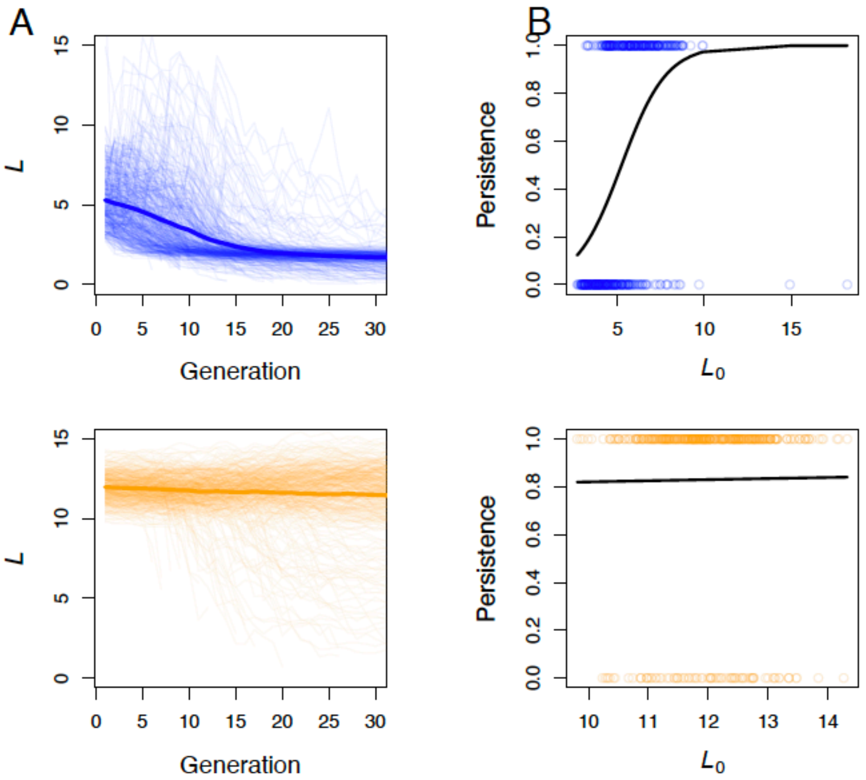


**Figure S24.** Temporal dynamics of the selection limit ($L$)(**A**), and the relationship between population viability and the selection limit at the beginning of the simulations ($L_0$) (**B**). Results are shown for simulated populations with a major locus responsible for 90% of $V_G$ (top row, blue lines), and where the selected trait was polygenic (bottom row, orange lines). The optimum phenotype in these simulations shifted from 100 to 110 over the first 10 generations, and the selected phenotype was plastic with plasticity parameter $m$ = 0.1 as described in the main text. The black lines in **B** are fitted logistic regression lines.

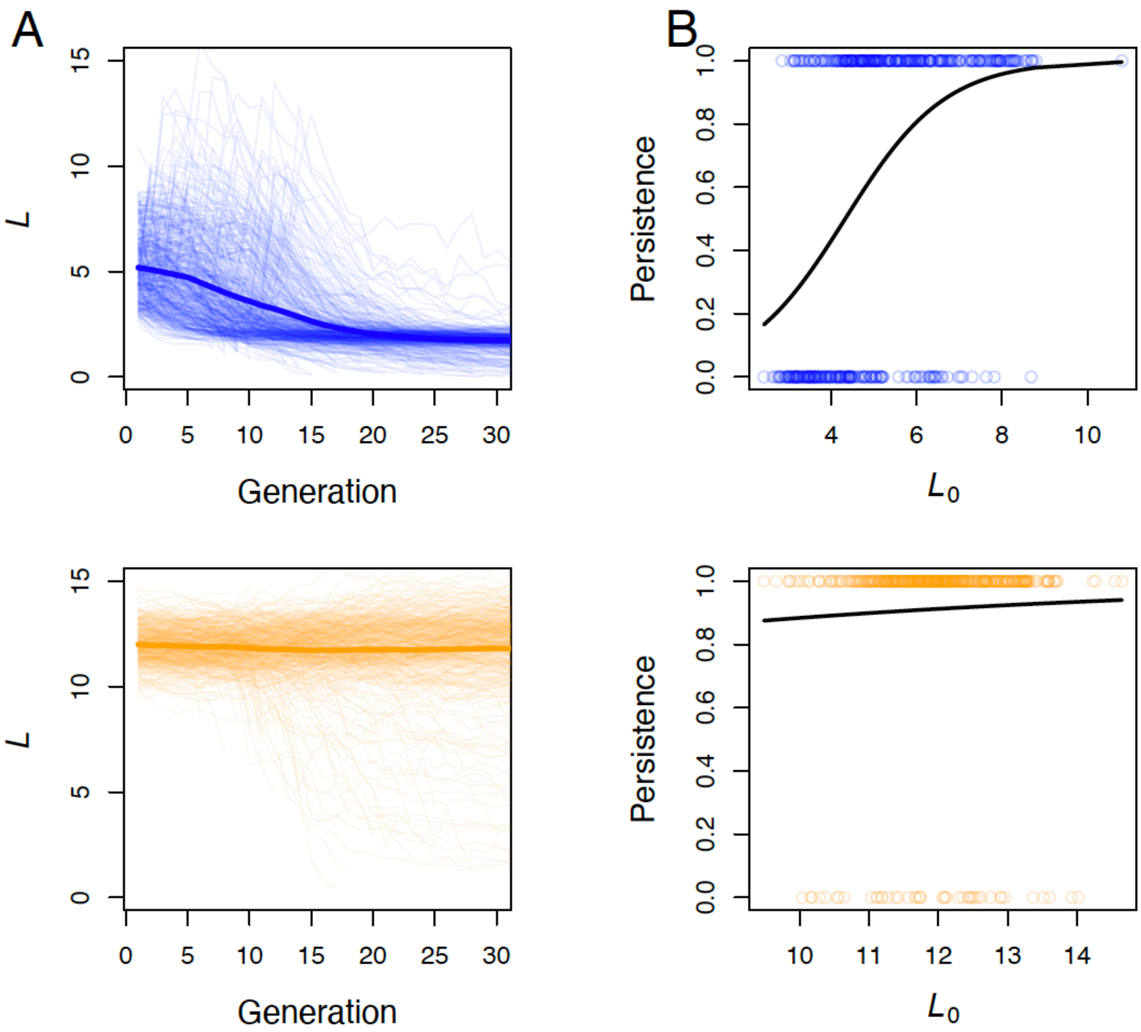


**Figure S25.** Temporal dynamics of the selection limit (*L*)(**A**), and the relationship between population viability and the selection limit at the beginning of the simulations ($L_0$) (**B**). Results are shown for simulated populations with a major locus responsible for 90% of $V_G$ (top row, blue lines), and where the selected trait was polygenic (bottom row, orange lines). The optimum phenotype in these simulations shifted from 100 to 110 over the first 10 generations, and the selected phenotype was plastic with plasticity parameter $m$ = 0.2 as described in the main text. The black lines in **B** are fitted logistic regression lines.

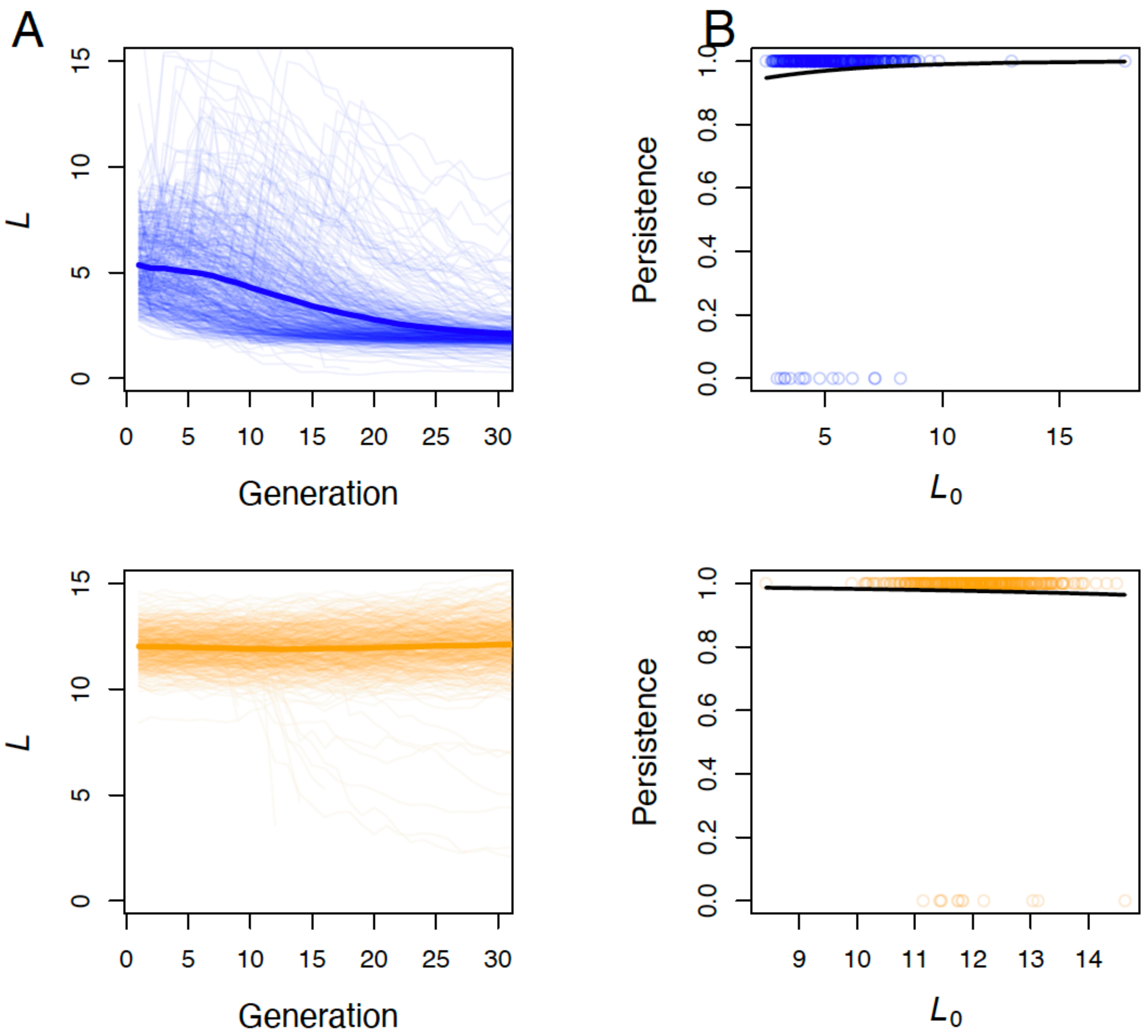


**Figure S26.** Temporal dynamics of the selection limit (*L*)(**A**), and the relationship between population viability and the selection limit at the beginning of the simulations ($L_0$) (**B**). Results are shown for simulated populations with a major locus responsible for 90% of $V_G$ (top row, blue lines), and where the selected trait was polygenic (bottom row, orange lines). The optimum phenotype in these simulations shifted from 100 to 110 over the first 10 generations, and the selected phenotype was plastic with plasticity parameter $m$ = 0.4 as described in the main text. The black lines in **B** are fitted logistic regression lines.

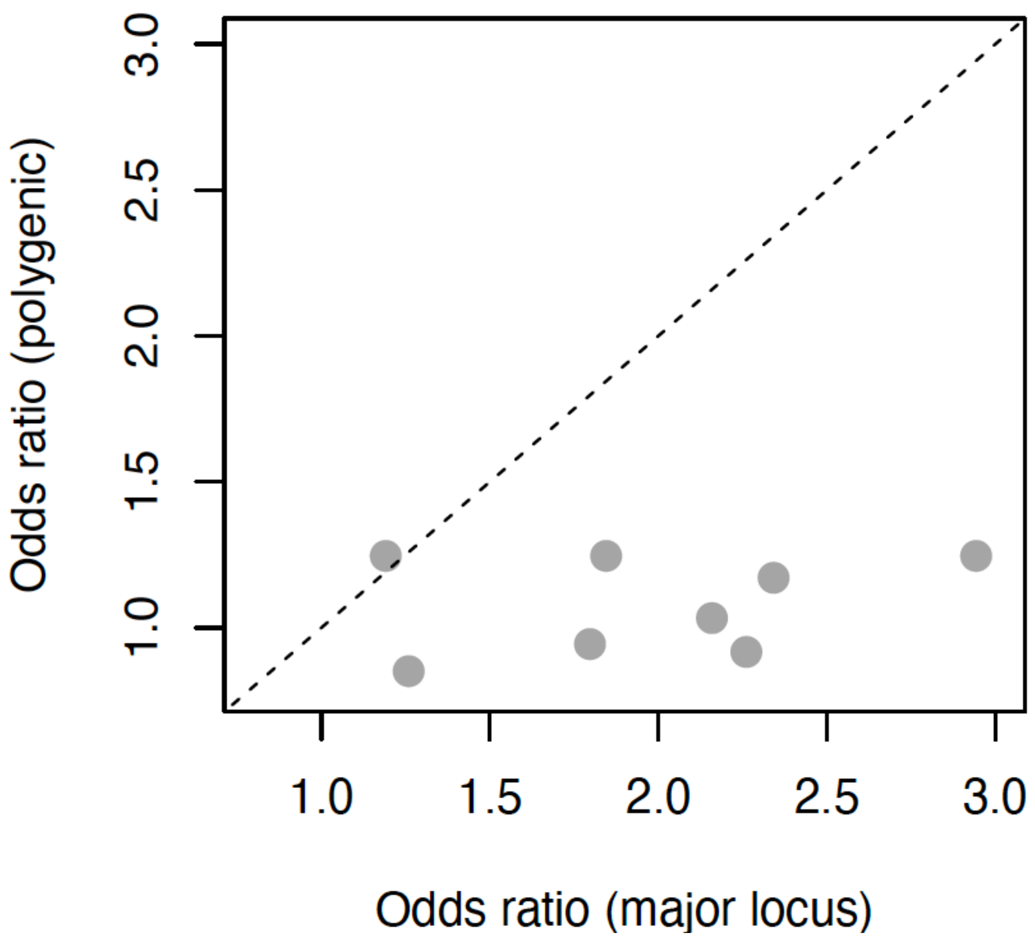


**Figure S27.** Odds ratios from logistic regressions of population persistence versus the short term selection limit ($L_0$) for simulations where the selected trait was polygenic (y-axis), and simulation scenarios with a large-effect locus (x-axis). The dashed line has an intercept and a slope of 1. Points below the line represent instances where the odds ratios from analysis of the polygenic scenario are smaller than the odds ratios from simulations with a large effect locus but were identical otherwise.

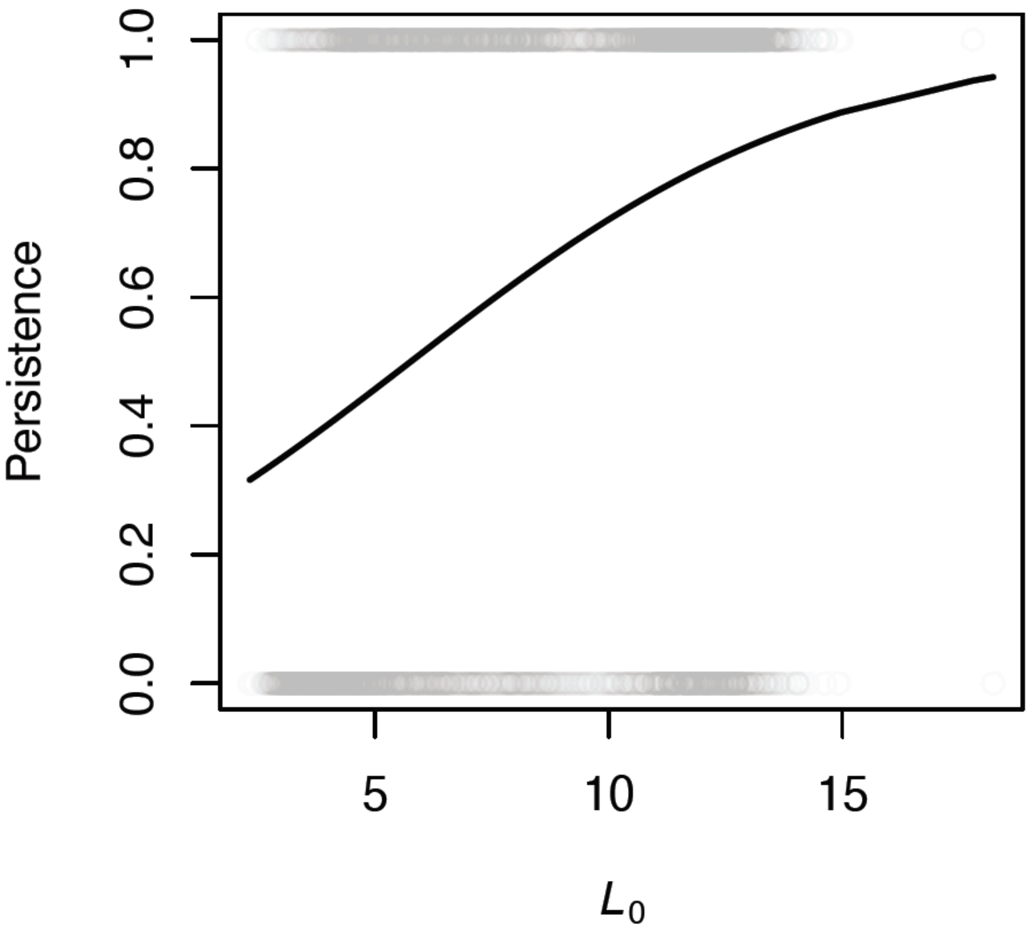


**Figure S28.** Population persistence (1 = population persisted to 80 generations, 0 = population went extinct within 80 generations) versus the initial short term selection limit ($L_0$). The fitted logistic regression line is shown in black.

**Table S1.** Logistic regression results from analyses of population persistence as a function of the initial short term selection limit ($L_0$) for simulations where the selected trait was governed in part by a large-effect locus. The data shown are the scenario that is being analyzed (scenario), the Figures where the simulation results are shown (Associated Figure(s)), the *P*-value from the regression analysis (*P*) and the associated regression coefficient estimate (*b*), and the odds ratio from the logistic regression model.

| Scenario | Associated Figure(s) | *P* | *b* | Odds Ratio (exp(b)) |
|---|---|---|---|---|
| $V_{QTL}/V_G$ = 0.9 | 3, S11 | 2.2x10-$^{29}$ | 1.08 | 2.943 |
| $V_{QTL}/V_G$ = 0.7 | S11 | 3.6x10$^{-15}$ | 0.612 | 1.845 |
| $V_{QTL}/V_G$ = 0.5 | S11 | 0.002 | 0.175 | 1.191 |
| 20 generations to theta1 | S13A | 1.8x10$^{-16}$ | 0.586 | 1.797 |
| 10 generations to theta1 | S13B | 1.8x10$^{-23}$ | 0.816 | 2.261 |
| plasticity (*m* = 0.1) | S16A | 1.7x10$^{-21}$ | 0.77 | 2.159 |
| plasticity (*m* = 0.2) | S16B | 2.9x10$^{-20}$ | 0.851 | 2.342 |
| plasticity (*m* = 0.4) | S16C | 0.219 | 0.23 | 1.259 |

**Table S2.** Logistic regression results from analyses of population persistence as a function of the initial short term selection limit ($L_0$) for simulations with a polygenic selected trait. The data shown are the scenario that is being analyzed (scenario), the Figures where the simulation results are shown (Associated Figure(s)), the *P*-value from the regression analysis (*P*) and the associated regression coefficient estimate (*b*), and the odds ratio from the logistic regression model.

| Scenario* | Associated Figure(s) | *P* | *b* | Odds Ratio (exp(*b*)) |
|---|---|---|---|---|
| Polygenic | 3, S11 | 0.058 | 0.22 | 1.25 |
| 20 generations to theta1 | S13A | 0.61 | -0.06 | 0.94 |
| 10 generations to theta1 | S13B | 0.48 | -0.09 | 0.91 |
| plasticity (*m* = 0.1) | S16A | 0.82 | 0.03 | 1.03 |
| plasticity (*m* = 0.2) | S16B | 0.39 | 0.16 | 1.17 |
| plasticity (*m* = 0.4) | S16C | 0.65 | -0.16 | 0.85 |
